# Developmentally regulated actin-microtubule crosstalk in Drosophila oogenesis

**DOI:** 10.1101/2024.12.12.628281

**Authors:** Wei-Chien Chou, Margot Lakonishok, Wen Lu, Vladimir I. Gelfand, Brooke M. McCartney

## Abstract

Actin-microtubules (MT) interactions are essential, but how those mechanisms are orchestrated in complex developing systems is poorly understood. Here we show that actin-MT crosstalk regulates actin cable assembly and the assembly and organization of MTs in Drosophila nurse cells during oogenesis. We found that a stable MT meshwork develops concurrently with actin cable initiation and requires acetylation for its maintenance. These γ-tubulin nucleated MTs are cortically tethered via Patronin and Shortstop, extend into the cytoplasm, and coalign with the elongating actin cables. The MT network promotes the cortical enrichment of the actin assembly factors Diaphanous and Enabled and loss of MTs dramatically reduced actin cable density and growth rate. We further demonstrated that actin filament assembly promotes cortical tethering of MTs and that loss of the actin filament bundler Villin/Quail results in fewer, shorter, and more highly coaligned MTs. Together, our data reveal multiple modes of coordinated actin-MT crosstalk producing actin cable-MT network synergy that is instrumental for oogenesis.

**Summary:** This study demonstrates actin–microtubule crosstalk in Drosophila oogenesis, revealing coregulation between actin filament assembly/bundling and microtubule nucleation/organization, highlighting the coordinated regulation of the cytoskeleton required during development.

## Introduction

Crosstalk between actin and microtubules (MTs) occurs across eukaryotes, with dynamic regulation between these two networks driving diverse cellular events, including cell migration, intracellular trafficking, and axon extension (Nicolas et al., 2009; Rodriguez et al., 2003). Actin-MT crosstalk can generally be attributed to a few core mechanisms, including crosslinking, shared regulators, and indirect effects arising from spatial and mechanical constraints (Dogterom & Koenderink, 2019; Pimm & Henty-Ridilla, 2021). For example, the single Drosophila spectraplakin, Short stop (Shot), directly crosslinks actin and MTs (Applewhite et al., 2013) and this dynamic linkage can promote the orientation of MT growth along actin bundles *in vitro* (López et al., 2014). Shared regulators, like the MT plus-end protein CLIP-170 (cytoplasmic linker protein-170) can recruit the formin mDia1 to assemble actin filaments on the plus ends of MTs *in vitro* (Henty-Ridilla et al., 2016). Other shared regulators have been identified in fission yeast, and in neuronal growth cones, fibroblasts, and focal adhesion sites in cultured cells (Chang & Martin, 2009; Dogterom & Koenderink, 2019; Juanes et al., 2019, 2020; Pimm & Henty-Ridilla, 2021; Wen et al., 2004). Spatial rearrangements of one network can influence the architecture of the other: actin bundle reorganization in growth cones can subsequently impact MT organization and dynamics (F.-Q. Zhou et al., 2002), and F-actin depletion in fibroblasts results in spatially unguided MT growth (Huda et al., 2012). Taken together, significant progress has been made in understanding actin-MT interactions, primarily through studies in minimal systems designed to pinpoint and characterize crosstalk mechanisms in relative isolation. The crosstalk between actin and MTs in intact tissues and organisms necessitates the coupling of various regulators and signaling events, temporal synchronization, and complex feedback loops (Pimm & Henty-Ridilla, 2021). The orchestration of these mechanisms *in vivo* is poorly understood.

Drosophila oogenesis has been a key model system for the identification of actin regulators and the investigation of developmentally regulated actin and MT functions (e.g. (Gates et al., 2009; Hudson & Cooley, 2014; Huelsmann et al., 2013; Logan et al., 2022; Lu et al., 2016; Roth-Johnson et al., 2014; Spracklen et al., 2014). Each *Drosophila* egg chamber, also known as a follicle, produces a single oocyte and comprises a syncytium of 16 germ cells connected by ring canals surrounded by a somatic epithelium of follicle cells (Fig. 1A1-2, S1B-F1). A posterior germ cell becomes the oocyte while the remaining 15 become nurse cells contribute mRNAs, proteins, and organelles to the growing oocyte (X. Wu et al., 2008). In a late stage of oogenesis (stage 11; s11), the nurse cells undergo cortical contractions to rapidly expel (“dump”) their remaining cytoplasm into the oocyte through the ring canals, roughly doubling its size in approximately 30 min (Fig. 1A3, S1G1). Prior to dumping, in s10B, dense arrays of actin cables initiate from the nurse cell cortex and elongate toward the nucleus, extending up to 30 µm (Fig. 1B4, D4). The actin cable arrays push and roll the nuclei away from the ring canals (Huelsmann et al., 2013; Logan et al., 2022). In mutants lacking actin cable arrays, the nuclei can obstruct the ring canals during dumping, blocking the completion of oogenesis (Cant et al., 1994; Gates et al., 2009; Logan et al., 2022; Mahajan-Miklos & Cooley, 1994).

**Figure 1:**
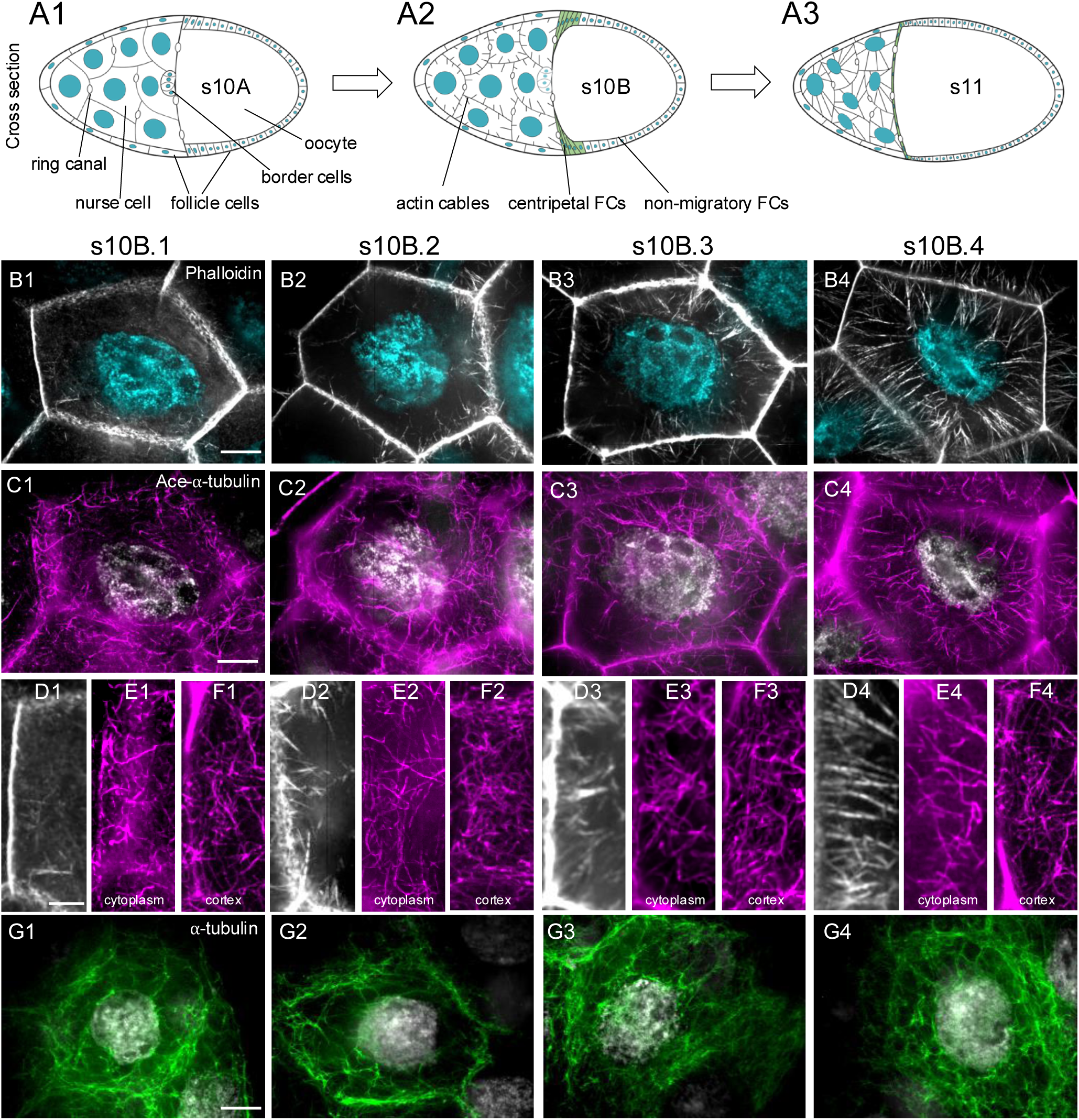
Wild type s10B Drosophila nurse cells contain a dense cortical and cytoplasmic MT network as actin cables assemble. (A1-A3) Schematic of stage 10-11 egg chambers. Images of fixed wild type stage 10B nurse cell actin cables stained with phalloidin shown in both low (B1-B4) and high (D1-D4) magnification. (C1-C4) Acetylated MTs (Ace-MTs) labeled with anti-acetylated-⍺-tubulin. High magnification of the cytoplasmic (E1-E4) and cortical (F1-F4) Ace-MT network. (G1-G4) Representative images of fixed wild type stage 10B nurse cells with total MTs labeled with anti-⍺-tubulin. Scale bars, 10µm (B,C,G) and 5µm (D,E,F).

**Figure 2:**
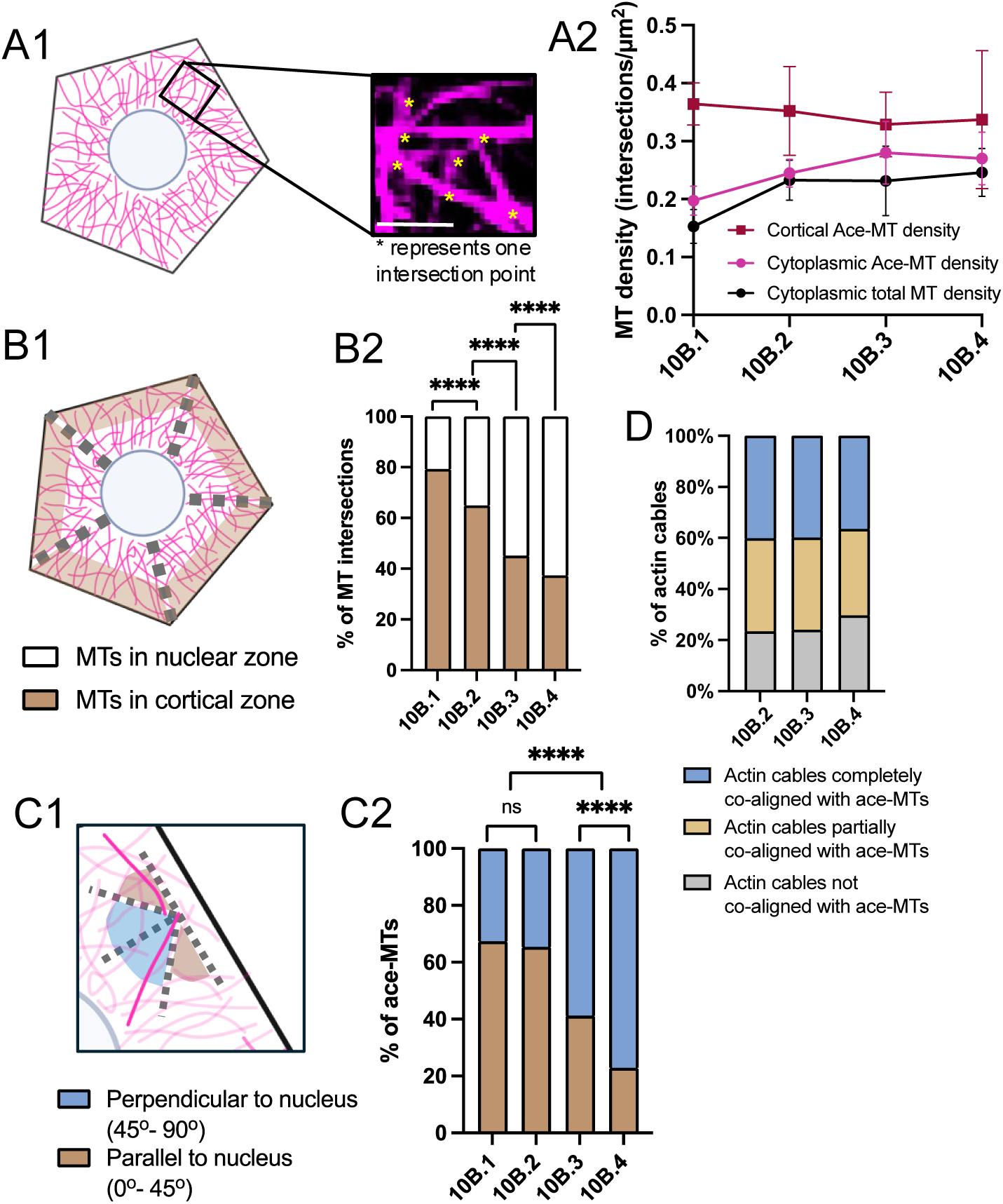
Quantification of features of the MT network in wild type s10B nurse cells. (A1-A2) Quantification of the density of the s10B total cytoplasmic MT network, and the cytoplasmic and cortical Ace-MT networks. Comparison across substages, one-way ANOVA, no significance, n=5-10 per substage, Tukey’s multiple comparison test. Comparison between total and acetylated cytoplasmic MT network density was not significant, two-way ANOVA, no significance, n=5-10 per substage. (B1-B2) Quantification of MT organization based on the density of cytoplasmic Ace-MTs in the cortical zone or the nuclear zone of the nurse cells. (C1-C2) Quantification of Ace-MT orientation relative to the nucleus. (B2,C2) Two-way ANOVA, n=5-10 per substage, ****p<0.0001, Tukey’s post-hoc analysis (D) Quantification of actin cable and Ace-MT co-alignment. No significance in the extent of co-alignment across different substages. Chi-square test for goodness fit, n=7-10 per substage, total=915 actin cables were analyzed.

The actin cables are composed of bundles of bundles of actin filaments mediated by Fascin (Singed) and Villin (Quail) (Cant et al., 1994; Guild et al., 1997; Matova et al., 1999). Each 2-4 μm unit bundle contains approximately 26 parallel actin filaments bundled in overlap to form mature cables (Guild et al., 1997). While cables initiate at roughly the same time at the start of s10B, their growth rates are regionally controlled by the differential localization of the actin assembly factors Diaphanous (Dia) and Enabled (Ena) (Logan et al., 2022). We further showed that this growth asymmetry is necessary for the directional repositioning of nurse cell nuclei prior to dumping (Logan et al., 2022). The mechanisms exerting temporal and spatial control over actin cable assembly upstream of the actin assembly factors Ena and Dia are not well understood. Prostaglandin (PG) signaling is required to produce normal actin cable arrays and can affect the cortical localization of Ena (Groen et al., 2012; Spracklen et al., 2014; Tootle & Spradling, 2008). PGs are known G-protein coupled receptor ligands, but the signaling cascade by which PGs regulate cytoskeletal organization is not known.

Here, we show that a dense cortical and cytoplasmic MT network plays an essential role in promoting the initiation and elongation of actin cables, and in turn, actin cable assembly is required to create and/or maintain the MT network. We found that the MT network in s10B nurse cells is composed of stable, acetylated MTs that co-align with actin cables. The cortex is a primary site for nucleating and organizing this acentriolar MT network, requiring γ-tubulin, CAMSAP/Patronin, and Spectraplakin/Shortstop. Reducing these proteins or MT acetylation significantly depleted both cortical and cytoplasmic MTs. Remarkably, depleting the MT network also caused a significant decrease in actin cable numbers and their elongation rate. Interestingly, direct loss of actin cables through reduction of actin filament assembly or bundling affected the MT network density, organization, orientation and actin-MT co-alignment in distinct ways. Our data support a model of dynamic cytoskeletal crosstalk in which a cortically nucleated and organized acetylated MT network is essential for actin cable assembly, and actin filament assembly and bundling are required to establish and maintain the MT network.

## Results

### Nurse cells contain dense, highly acetylated microtubule networks that coalign with growing actin cables

Nurse cell actin cable assembly occurs in s10B of oogenesis that is also defined by the centripetal migration of somatic follicle cells between the oocyte and the nurse cells (Fig. 1B). The developmental time points within s10B have been typically categorized as early, mid, and late. To increase the temporal resolution of actin cable assembly, we defined four 10B substages in fixed tissue based on two somatic cell populations: the migration of centripetal follicle cells and the visualization of border cells (Fig. S1 and Table.1). Because our genetic manipulations primarily target the germline, the wild-type somatic cells are unaffected developmental landmarks. This approach allows for accurate, comparative assessment of cytoskeletal development in germline knockdowns and improves the temporal resolution of the cable assembly process. Detailed descriptions of each stage are included in Table.1 and Fig. S1. While we cannot rule out potential indirect effects of the manipulated germline on the somatic cells, we have found this to be a tractable and reproducible staging method. In wild-type nurse cells, cables initiate from the cortex at s10B.1 (Fig.1B1, D1) and appear to elongate at a constant rate throughout s10B (Fig. S1I), consistent with what we have observed using live imaging (Logan et al., 2022). A robust actin cable array formed by s10B.3, with most cables reaching roughly halfway toward the nucleus (Fig.1B3, D3). In s10B.4, cable density peaks at 0.5 cables/µm, and length reaches 13 µm (Fig.1B4, D4, S1H-I). Some cables have contacted or are nearing nuclei in preparation for repositioning and s11 dumping (Fig. S1G5).

The MT network in nurse cells is challenging to visualize: MTs cannot be resolved with GFP-tagged alpha-tubulin (Logan & McCartney, 2020) and MTs are difficult to fix and label with anti-tubulin antibodies (Legent et al., 2015). An anti-acetylated tubulin antibody revealed a robust meshwork of cytoplasmic MTs throughout s10B (Fig. 1C1-4, E1-4). To ask if the acetylated MTs (Ace-MTs) represent all or a subset of MTs in the network, we labeled the total MT network following cytoplasmic extraction (adapted from (Lu et al., 2022)). This process disrupts the cortex and actin cables but allows us to visualize the total cytoplasmic MT network. We found that the total cytoplasmic MT density did not significantly differ from the cytoplasmic Ace-MT density (Fig. 2A2). Because labeling the Ace-MTs allowed us also to preserve the cortex and cables, we used that as our primary marker for the MT network.

Overall, Ace-MTs form an intertwined cytoplasmic network that appeared to extend from a dense cortical meshwork (Fig. 1F1-4). Consistent with this, we observed MTs with one end embedded in the cortex (Fig. S3A1-3) or lying parallel to the cortex with the other end extending outward into the cytoplasm (Fig. S3B). Occasionally, we observed prominent perinuclear rings where MTs appeared to lie parallel to or circumnavigate the nucleus (Fig. S3D1-3). In pre-stage 10B egg chambers, MTs glide through ring canals as part of the transport system into the oocyte (Lu et al., 2022; Moon & Hazelrigg, 2004; Nicolas et al., 2009). We also observed cases of MTs in nurse cell ring canals in s10B (Fig. S3C1-2).

Because the network is generally too intertwined to identify and count individual MTs, we quantified MT network density by counting the number of Ace-MT intersections per µm^2^ (Fig. 2A1 and S2A1-2). We found that unlike the increasing density of actin cables throughout s10B (Fig. S2A), the density of the cytoplasmic and cortical Ace-MT networks did not change significantly. Because there is variation in cable and Ace-MT density between egg chambers, we asked if these correlated but found no relationship (Fig. S1J). Although there were no significant changes in Ace-MT network density, the organization and orientation of the Ace-MT network changed through s10B at the same time actin cables elongate toward the nuclei. A significant shift in Ace-MTs organization was observed from s10B.1 to s10B.4 (Fig. 2B2), with MTs transitioning from higher density near the cortex (referred to as the ’cortical zone,’ shaded brown in Fig. 2B1) to higher density near the nucleus (referred to as the ’nuclear zone,’ unshaded in Fig. 2B1) (Fig. S2B). In the orientation of Ace-MTs relative to the nucleus in s10B.1 to s10B.2, approximately 65% were oriented roughly parallel to the cortex (0-45°; brown shaded area, Fig. 2C1), while 35% were oriented roughly toward the nucleus (46-90°; termed “nuclear oriented”, blue shaded area, Fig. 2C1) (Fig. S2C1-3). This pattern shifts markedly in s10B.3 to 10B.4, where the majority of Ace-MTs orient toward the nucleus (Fig. 2C2). Interestingly, this coincides with the period of time when actin cables are nearing or contacting the nuclei, suggesting a potential relationship between MT orientation and the elongation of the cable network. We quantified actin cable-MT coalignment through s10B and found that over 80% of actin cables exhibited some degree of coalignment with Ace-MTs and that did not change over time (Fig. 2D, Fig. S2D1-F3). Such frequent co-alignment suggests the potential for functional interactions.

### The microtubule network exhibits a developmentally regulated enhancement of stability at stage 10B

Acetylated tubulin is a well-established indicator of MT stability; whether acetylation is the cause or consequence of stability, or both, is not fully understood (Janke & Magiera, 2020). Labeling the total MT network with Tubulin-Tracker enables visualization in live egg chambers (Logan & McCartney, 2020), but it is challenging to find MT ends in the dense, intertwined network, making it difficult to assess MT dynamics directly. Furthermore, there are stabilizing effects of such taxol-derived fluorescent probes (Logan & McCartney, 2020; Lukinavičius et al., 2014). Therefore, we took two alternative approaches to investigate the dynamic properties of the cytoplasmic MT network.

First, we tracked growing MT plus ends with GFP-tagged End-Binding protein 1 (EB1), a plus-end tracking protein (+TIP), expressed under a ubiquitous promoter (*ubi-EB1-GFP*) (Lu et al., 2023; Vaughan, 2005). We imaged egg chambers every 2 sec over a period of 2 min and quantified MT dynamics by assessing the frequency of EB1 “comets”, indicators of growing MT plus ends (Lu et al., 2021). We classified egg chambers as having frequent comets if >30% of the frames contained at least one nurse cell with at least one EB1 comet, while those with few comets had <30% of frames with EB1 comets. There was a striking difference between MT dynamics in s10B nurse cells compared to earlier nurse cells or follicle cells at any stage. Consistent with previous reports (Lu et al., 2022), nurse cells prior to s10B exhibited abundant EB1 comets running parallel to the cortex and throughout the cytoplasm (Fig. 3A1, supplemental video 1). As the nurse cells transitioned into s10B and the onset of cable assembly, the frequency of EB1 comets dropped dramatically (Fig. 3A2, supplemental video 3). Whereas at s9 and s10A 100% of the egg chambers had frequent comets (supplemental video 2), by s10B.1 that had dropped to roughly 30% and by s10B.4 EB1 comets were rarely seen (Fig. 3B). When comets were observed in s10B, they were predominantly near and parallel to the cortex (supplemental video 3).

**Figure 3:**
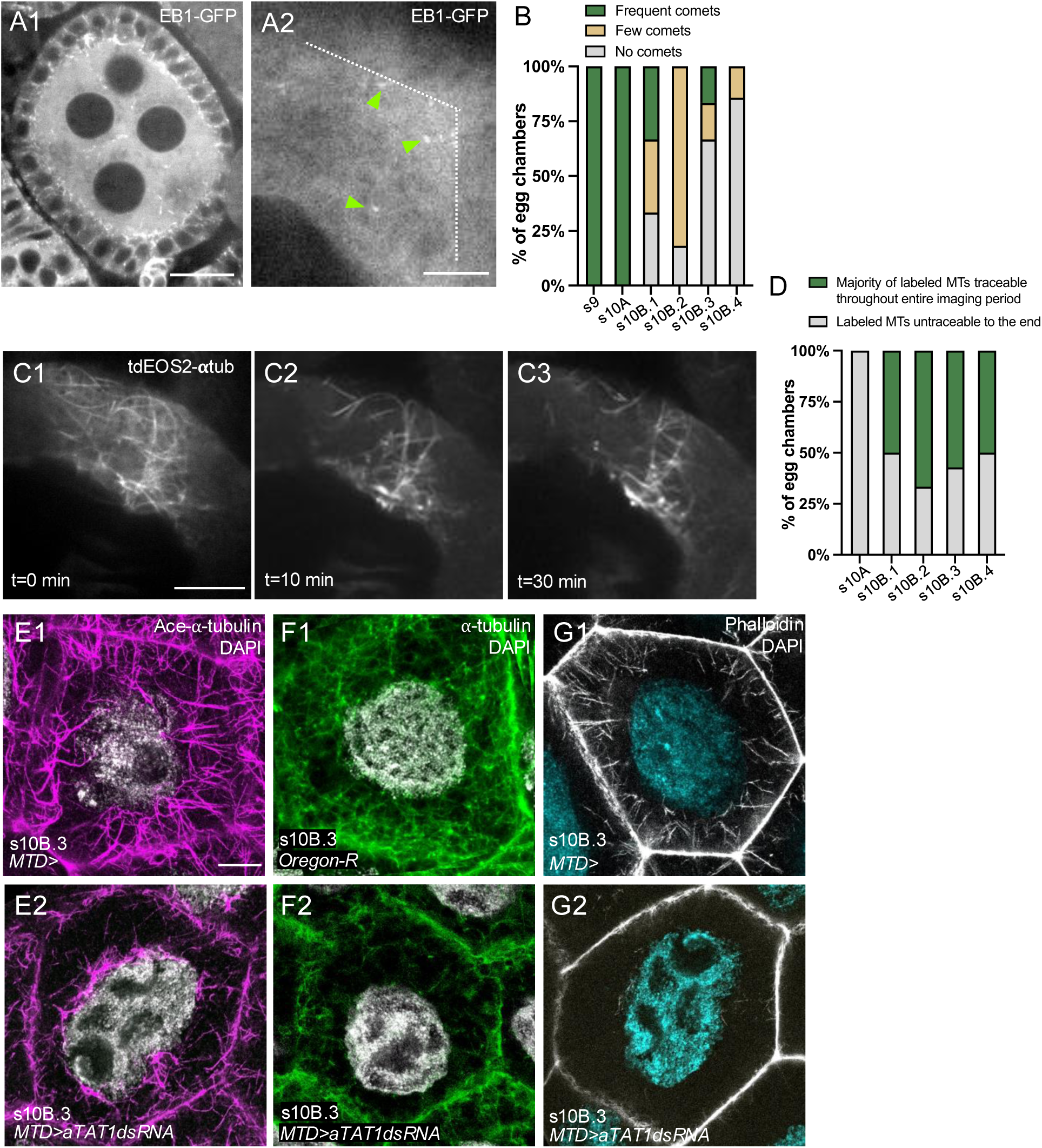
The nurse cell Ace-MT network switches from more dynamic to more stable at s10B. (A1-A2) Live imaging of EB1-GFP to track growing MT plus ends. (A1) In s2-3, EB1 comets were abundant, distributed throughout the cytoplasm, and parallel to the cortex. Scale bar, 20µm. (A2) In s10B.2, EB1 comets were sparse (comets pointed with green arrowheads, cortex marked by white dotted line). Scale bar, 5µm. (B) Categorization of egg chambers based on EB1 comet frequency from s9 to s10B.4. Egg chambers were classified as having “frequent comets” if >30% of frames contained at least one nurse cell with an EB1 comet, as having “few comets” if <30% of frames showed EB1 comets. (See also supplemental videos 1, 2). (C1-C3) Live imaging of a portion of the MT network in s10B.2 nurse cells using locally photoconverted tdEOS2-αtub. A subpopulation of MTs at time zero post-photoconversion (C1) remained stable over 30 minutes (C2,C3). Scale bar, 10 µm. (D) Categorization of photoconverted egg chambers from s10A to s10B.4 based on converted-MTs traceability. Egg chambers were classified as having labeled MTs where the majority could be tracked until the end of the imaging period or as having labeled MTs where the majority could not be tracked for both 10-minute and 30-minute imaging durations. (E-G) aTAT1 is required for the acetylation and maintenance of s10B nurse cell MT network and for actin cable assembly. Representative images of the s10B.3 (E1-2) ace-MT and (F1-2) total MT network in control (*MTD>*) wild-type and *MTD>aTAT1 dsRNA*. Reducing aTAT1 acetylation significantly reduced the (E2) acetylated and (F2) total cytoplasmic MT network. (G1-2) Depletion of the MT network resulted in a dramatic depletion of actin cables. Scale bar, 10 µm. (B,D) n=4-12 egg chambers per sub/stage.

As a complementary approach, we used photoconversion of α-tubulin tagged with a photoconvertible protein, tdEOS, to label and track a subpopulation of MTs in live nurse cells. This overcame the challenges of reduced resolution due to egg chamber thickness and high cytosolic tubulin levels in nurse cells (Lu et al., 2013, 2016, 2022). We photoconverted a region of the cytoplasmic MT network and tracked the labeled MTs every 10 sec for 10 minutes or 30 sec for 30 min. In all cases, uniformly labeled MTs persisted to the end of the imaging period (Fig. 3C1-3, D, supplemental video 4). Because MTs can move relative to each other and out of the imaging plane, we classified nurse cells as either having labeled MTs where the majority could be tracked to the end of the imaging period or having labeled MTs where the majority could not be tracked to the end of the imaging period. Regardless of whether we imaged for 10 or 30 min, at least 50% of egg chambers throughout s10B had uniformly and persistently labeled MTs that could be tracked to the end of the imaging period (Fig. 3D). Together, these results suggest that there is a developmentally regulated shift toward enhanced stability of the MT network in nurse cells coincident with the onset of actin cable assembly.

The s9 MT network is characterized by frequent EB1 comets, both near the cortex and through the cytoplasm (supplemental video 2; (Lu et al., 2021)). Consistent with that, we found that the s9 cytoplasmic MTs are typically non-acetylated though we did observe acetylated MTs at the cortex (Fig. S3E1-2). To ask if acetylation is required for the stability of the s10B MT network, we reduced the level of the major Drosophila ⍺-tubulin acetylase (aTAT1) by driving *aTAT1* dsRNA expression with a strong germline driver (MTD>; germline driver). Reducing aTAT1 levels led to a significant loss of both cortical and cytoplasmic acetylated MTs in s10B (Fig 3E1-2, 6E-F). Remarkably, aTAT1 reduction also caused a dramatic loss of total cytoplasmic MTs (Fig. 3F1-2) suggesting the acetylation that increases significantly in s10B (Fig. S3E1-G2) is required for the overall maintenance of the entire s10B MT network.

### The nurse cell cortex is the primary ncMTOC

Like many differentiated cells, the nurse cell MTOC is not the centrosome at s10B; all nurse cell centrosomes migrate into the oocyte at s2 (Bolívar et al., 2001), and no nurse cell MTOC has been identified after s2. A growing number of non-centrosomal MTOCs (ncMTOCs) have been identified in a wide variety of cell types in mammals and Drosophila (Sanchez & Feldman, 2017). In some Drosophila neurons, γ-tubulin localizes to the Golgi, which serves as an MTOC (Mukherjee et al., 2020). By s10B, the nurse cell Golgi have been transported into the oocyte (Bastock & Johnston, 2008). In Drosophila follicle cells, a ncMTOC is assembled at the apical cortex, and in s9 oocytes, the primary ncMTOC is the anterior cell cortex (Nashchekin et al., 2016). During both mammalian and Drosophila muscle development, MTOCs relocate from the centrosomes to the nuclear envelope (Starr, 2017). The Drosophila larval fat body cells also assemble ncMTOCs on the nuclear envelope (Zheng et al., 2020). Given the distribution of MTs we observed in s10B nurse cells, we predicted that the cortex is the primary ncMTOC. To test this, we depolymerized MTs using nocodazole and assessed Ace-MT shrinkage to their origins in *ex vivo* culture (Groen & Tootle, 2015). After 30 min of treatment, we found a substantial depletion of cytoplasmic Ace-MTs compared to the DMSO control (Fig. 4A1-2, B1-2, D). After 60 min of treatment, only a small number of cytoplasmic Ace-MTs remained (Fig. 4C1-2, D). Consistent with a collapse of the Ace-MT network towards the cortex, after 30 or 60 min of treatment there was a significant reduction in Ace-MT density within the nuclear zone, with the remaining MTs primarily in the cortical zone (Fig 4E). At the cortex, Ace-MT density also significantly decreased after nocodazole treatment (Fig 4A3-C3), but at a slower rate than in the cytoplasm (Fig 4D) suggesting that the cortical network may be more resistant to nocodazole. We observed occasional perinuclear MTs in wild type nurse cells (Fig. S4D1-2), raising the possibility of an additional nuclear envelope ncMTOC. While we can’t rule that out, these results suggest that the cortex is the primary ncMTOC in s10B nurse cells.

**Figure 4:**
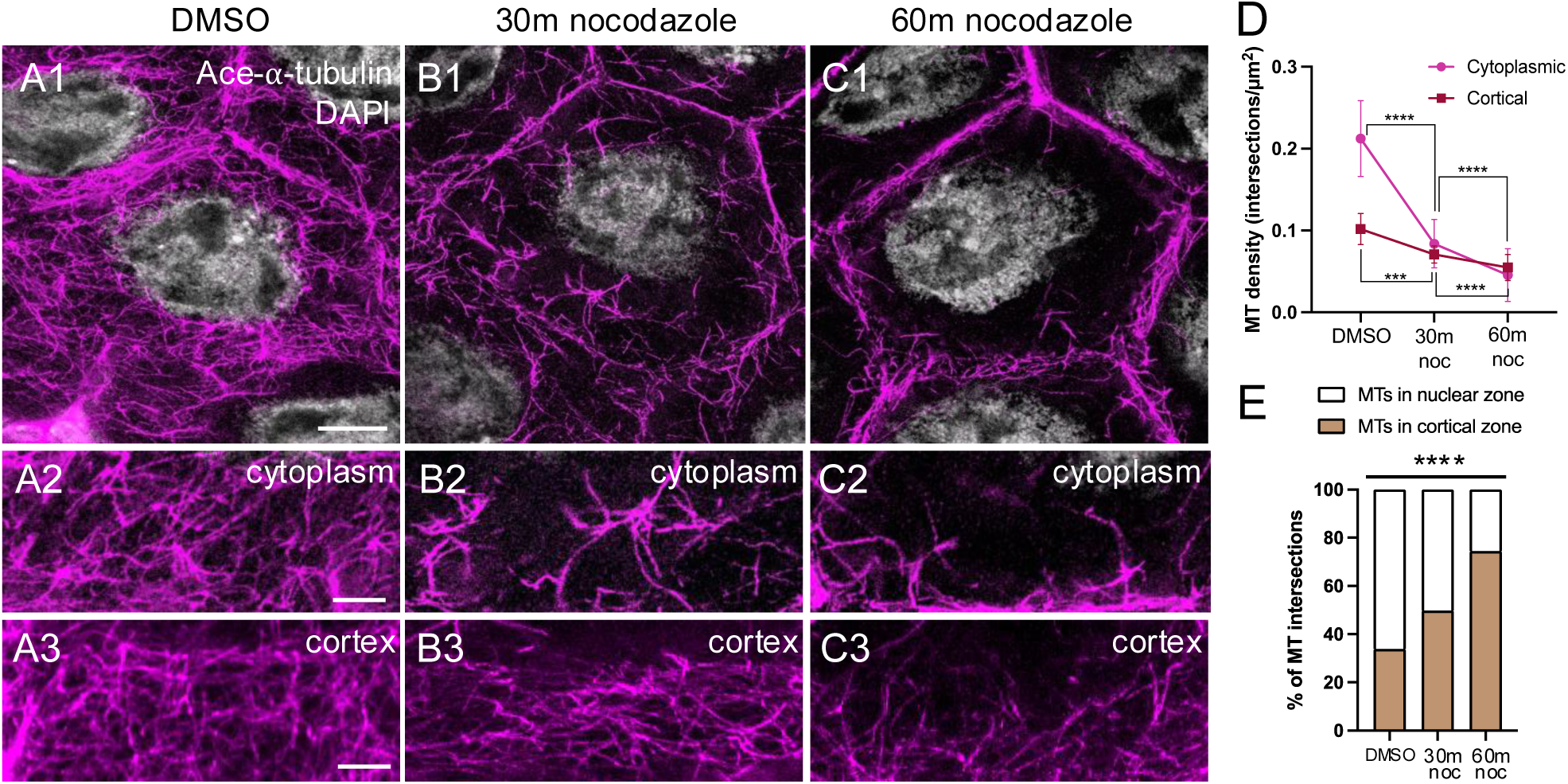
Nocodazole depolymerization shrank the Ace-MT network back to the cortex. (A1-A3) The Ace-MT network was unaffected by 60 minutes of DMSO treatment. (B1-B3) After 30 minutes of nocodazole treatment, there was a substantial depletion of cytoplasmic Ace-MTs and a reduction in cortical Ace-MTs. (C1-C3) After 60 minutes of nocodazole treatment, very few cytoplasmic Ace-MTs remained, and there was a significant reduction in cortical Ace-MTs. Scale bar, 10 µm (A1-C1); Scale bar, 5 µm (A2-C2, A3-C3). (D) Cytoplasmic and cortical Ace-MT density after 30 or 60 min nocodazole treatment or 60 min of DMSO treatment. (E) Cytoplasmic Ace-MT organization showed a significant shift toward the cortical zone as Ace-MTs in the cytoplasmic zone depolymerized after nocodazole treatment. (D, E) One-way ANOVA, ****p<0.0001, n=8-11 per substage, Tukey’s multiple comparison test.

### *γ*-tubulin, Patronin and Shot are required to produce the nurse cell MT network

ncMTOCs employ MT nucleators, as well as stabilizers and anchors—proteins associated with MT minus ends and proteins that couple these minus-end proteins to specific subcellular locations (Sanchez & Feldman, 2017). In the s9 Drosophila oocyte that employs a cortical ncMTOC, *γ*-tubulin is the likely nucleator, the CAMSAP Patronin is the stabilizer, and the spectraplakin Shortstop (Shot) anchors Patronin, and thus MTs, to the cortex (Nashchekin et al., 2016). This triad is conserved in other systems (Sanchez & Feldman, 2017; Tillery et al., 2018) and we predicted that it may function as the s10B nurse cell ncMTOC.

Anti-*γ*-tubulin clearly labeled centrosomes in early-stage egg chambers (Fig. S4C3, yellow arrowheads). Surprisingly, *γ*-tubulin was not significantly enriched at any subcellular location in s10B nurse cells, though we did observe occasional, weak cortical or perinuclear enrichment (Fig. S4A1-C2, yellow arrowheads). Labeling *γ*-tubulin after cytoplasmic extraction to enhance visualization of a potential membrane-associated pool (Lu et al., 2022) yielded the same results (data not shown). In contrast, both Patronin and Shot were strongly enriched at the cortex (Fig. 5A1-2, S4D1-2). In addition, Shot localized to actin cables (Fig. 5B1-2) and Patronin was occasionally enriched in the cytoplasm near the nucleus (Fig. S4E1-2). Consistent with the model that Shot tethers Patronin to the cortex as in s9 oocytes, Patronin’s cortical localization was lost in Shot knockdown but not vice versa (Fig. S4G1-2). We reduced *γ*-tubulin, Patronin, or Shot in nurse cells using dsRNA expression: **γ*-tubulin23C* and *Patronin* dsRNAs were driven with a strong germline driver (MTD>), but because strong Shot reduction blocked oogenesis prior to s10B (Lee et al., 2016; Lu et al., 2021) *shot* dsRNA was expressed using the weaker MDD-GAL4 (MDD>; germline driver). All resulted in significant loss of the cortical and cytoplasmic Ace-MT networks consistent with a role in cortical ncMTOC activity (Fig. 6A2-D2, A4-D5, E, F).

**Figure 5:**
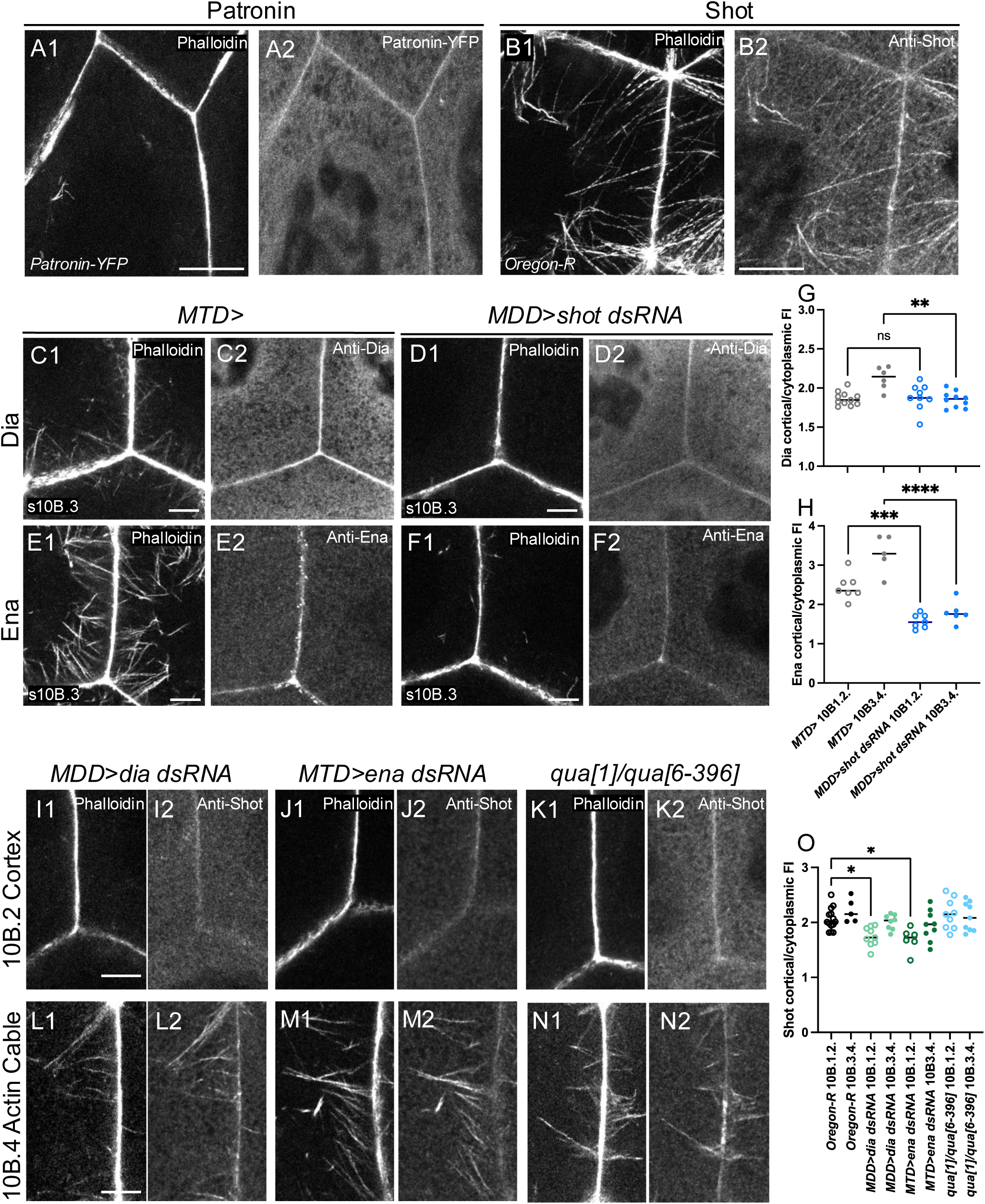
Patronin and Shot are enriched at the cortex. Shot reduction disrupted the cortical localization of Dia and Ena and vice versa. (A1-A2) Patronin enrichment at the cortex visualized by endogenously tagged Patronin-YFP in fixed s10B.1 nurse cells. (B1-B2) Wild type Shot was enriched at the cortex and localized to actin cables in s10B.4. Scale bars, 10 µm. Dia (C1-C2) and Ena (E1-E2) were enriched at the cortex in control (*MTD>*) s10B.3 nurse cells but that enrichment was lost in Shot knockdowns (D1-D2; F1-F2; *MDD>shot dsRNA*). Quantification of Dia (G) and Ena (H) cortical enrichment. s10B.1 and s10B.2 were grouped and s10B3 and s10B.4 were grouped. (I1-N2) Shot localization to the cortex (I1-K2) or to actin cables (L1-N2) in Dia or Ena knockdowns, or Villin/Quail mutant nurse cells. Shot’s cortical enrichment, but not actin cable enrichment, was reduced in Dia or Ena knockdowns. (O) Quantification of Shot cortical enrichment. s10B.1 and s10B.2 were grouped and s10B3 and s10B.4 were grouped. Scale bars, 5 µm. (G,H,O) One-way ANOVA, *p<0.05, **p<0.01, ***p<0.001, ****p<0.0001, n=as plotted. Tukey’s multiple comparison test.

**Figure 6:**
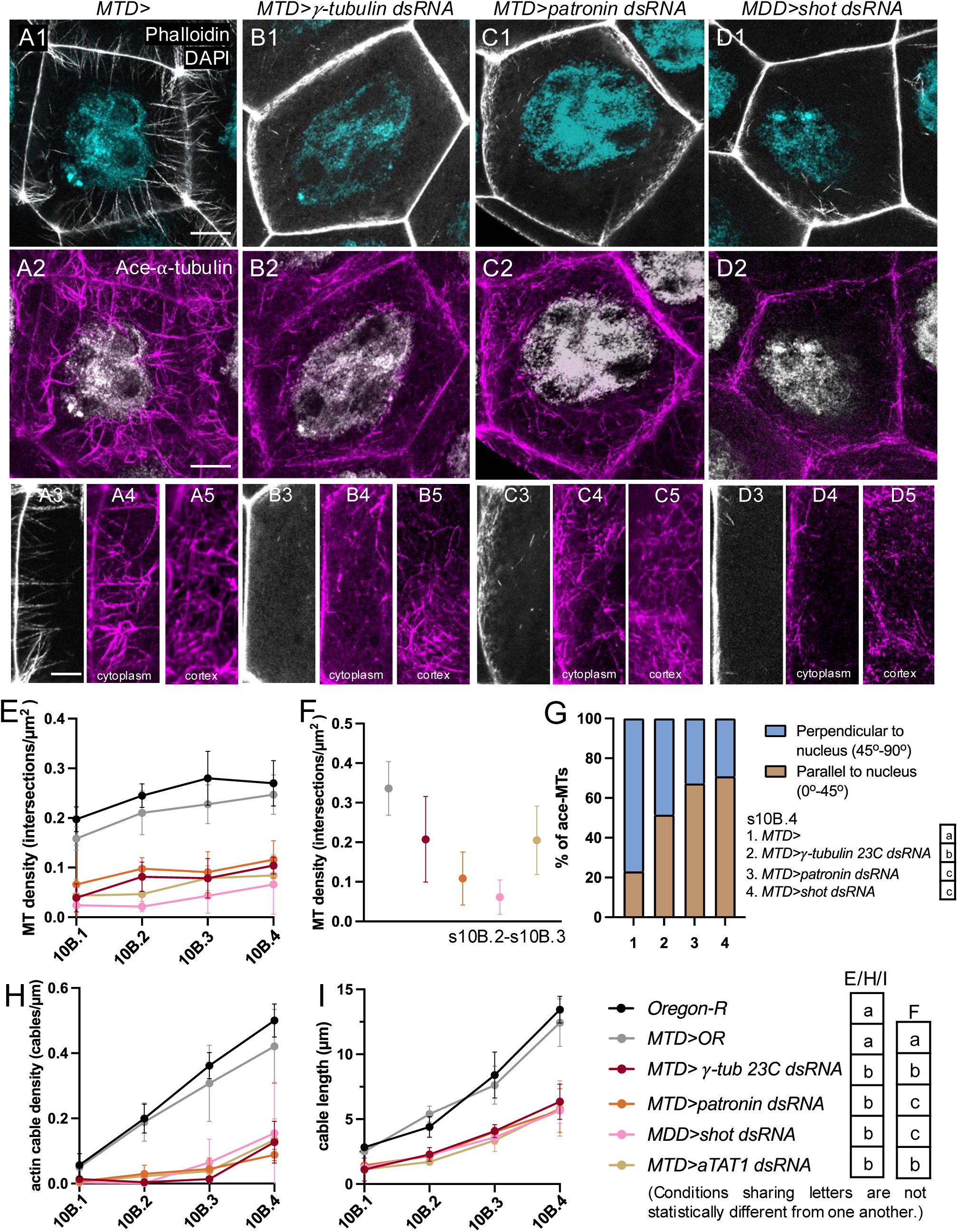
The nurse cell Ace-MT network is required for actin cable assembly. (A1-D5) Representative fixed images of s10B.3 actin cables and Ace-MT network in control *MTD>* (A1-A5), *MTD>*γ*-tubulin dsRNA* (B1-B5), *MTD>patronin dsRNA* (C1-C5) and *MDD>shot dsRNA* (D1-D5) nurse cells. (A1-D2) Scale bars, 10 µm; (A3-D5) Scale bars, 5 µm. (E) Quantification of cytoplasmic and (F) cortical Ace-MT density upon depletion of γ-tubulin, Patronin, Shot, or aTAT1. (G) Quantification of Ace-MT orientation for s10B.4. γ-tubulin, Patronin, Shot or knockdowns increased the fraction of Ace-MTs oriented parallel to the nucleus. Loss of Ace-MTs significantly reduced actin cable density (H) and actin cable length (I) across s10B. (E, F, H, I) Share figure legend. Two-way (E, G, H, I) or one-way (F) ANOVA with Tukey’s multiple comparison test. Conditions sharing letters are not statistically different from one another.

Reduction of either *γ*-tubulin 23C or Patronin produced very similar decreases in cortical and cytoplasmic Ace-MT density (Fig. 6E). While fewer MTs were oriented towards the nucleus at s10B.4 in those knockdowns (Fig. 6G), the depleted cortical and cytoplasmic Ace-MTs networks retained a similar morphology to those in wild type nurse cells: a MT mesh with relatively long individual MTs (Fig. 6B2,4,5, 6C2,4,5). Reduction of Shot produced a significantly stronger loss of both the cortical and cytoplasmic Ace-MT networks (Fig. 6D2,4,5, E, F). Interestingly, only short fragments and Ace-MT stumps remained in the Shot knockdown (Fig. 6D2,4,5), consistent with MT nucleation, but lack of elongation and/or stabilization. If Shot’s role is solely to cortically tether Patronin and MT minus ends, we predicted that some long MTs extending from the cortex would remain in the knockdown, similar to the Patronin knockdown. Their absence, but the presence of Ace-MT stumps, suggests that Shot may have additional roles, though we can’t rule out the possibility that differences in knockdown efficiency are the cause of this distinction. Given Shot’s localization to actin cables (Fig. 5B1-2) and its known molecular functions (Lu et al., 2021; Nashchekin et al., 2016; Sun et al., 2019), crosslinking the actin cables and the co-aligning cytoplasmic Ace-MTs is a possible additional role. Regardless, these data support the model that the nurse cell Ace-MT network is organized by a cortical ncMTOC that requires γ-tubulin, and Patronin and Shot.

### Microtubules are required for actin cable assembly

Remarkably, dramatic depletion of the MT network through ⍺TAT1, γ-tubulin, Patronin, or Shot reduction led to a similarly dramatic depletion of actin cables (Fig. 3G1-2, 6A1-D1, H). We found a nearly complete absence of cable initiation in s10B.1 and s10B.2 and the presence of only a few, short cables in s10B.3 and s10B.4 (Fig. 3G1-2, 6A1-D1, H). The remaining cables showed a significantly reduced growth rate compared to wild type (Fig. 6I). These defects in actin cable density and growth rate were comparable to what we observed when we directly disrupted actin cable assembly through the reduction of Dia, Ena, or Villin/Quail (Fig. 7A1-D1, E, F). Because the cortex harbors an extensive MT meshwork and is a known site of actin filament and cable assembly, we asked if MTs are required for Ena and Dia’s cortical localization in nurse cells using *MDD>shot dsRNA* that produced some of the strongest MT depletions. In s10B.1 and s10B.2 when cable assembly is initiating in wild type nurse cells, we observed a significant reduction in Ena’s cortical enrichment while Dia’s localization was unaffected (Fig. 5G, H). By s10B.3 and s10B.4, both Ena and Dia were significantly less enriched at the cortex (Fig. 5C1-F2, G, H). Thus, the reduction in cortical Ena and Dia could contribute to the loss of actin cables in MT depleted nurse cells. However, the fact that there is virtually no cable initiation in s10B.1 and s10B.2 when Dia remains cortically enriched and Ena is only moderately reduced strongly suggests that there will be other roles for MTs as well.

**Figure 7:**
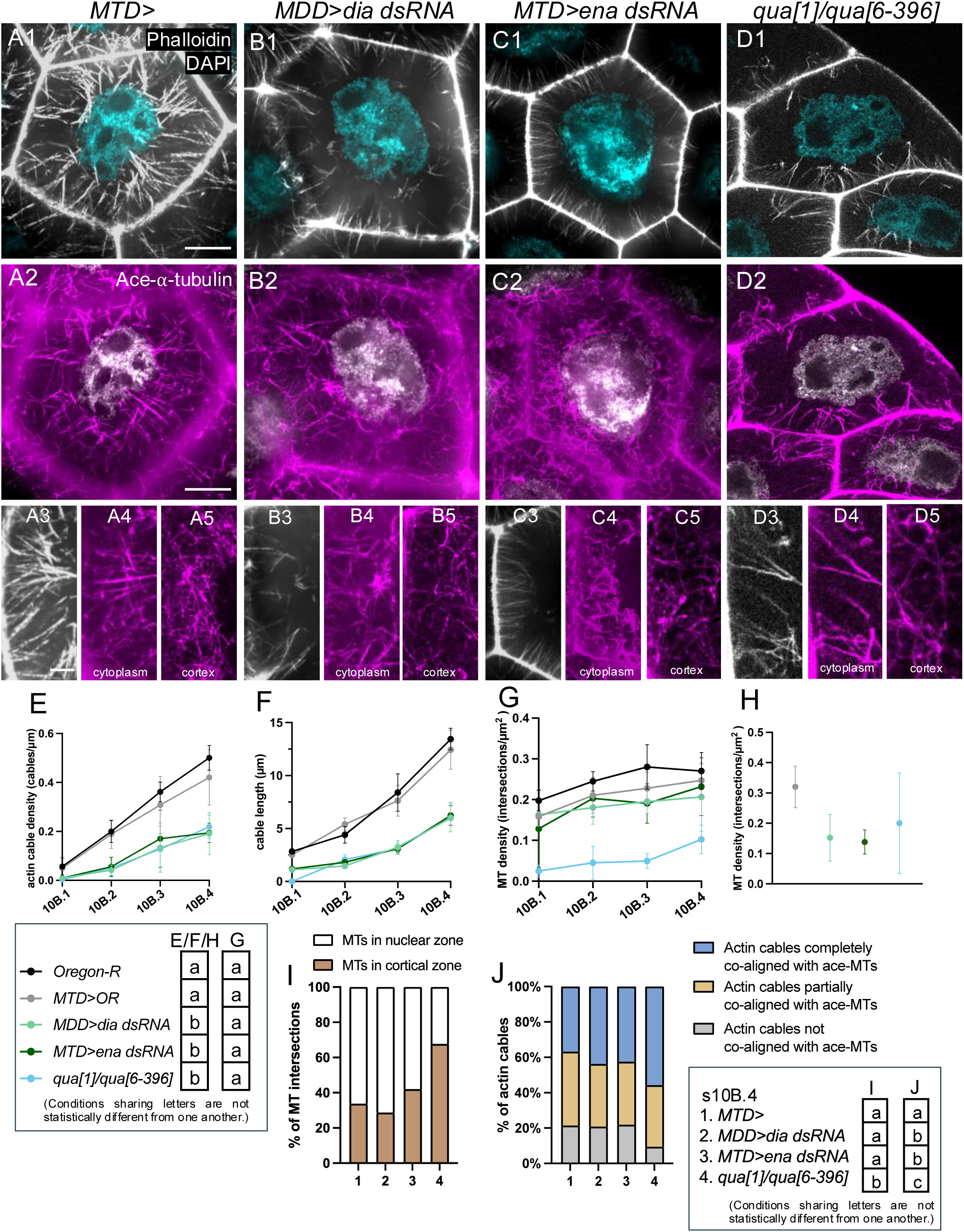
Direct perturbation of actin filament assembly and bundling led to comparable defects in actin cable assembly, but varied Ace-MT network defects. (A1-D5) Representative fixed images of s10B.4 actin cables and Ace-MT network in control (A1-A5), *MDD>dia dsRNA* (B1-B5), *MTD>ena dsRNA* (C1-C5) and *qua[1]/qua[6-396]* (D1-D5) nurse cells. (A1-D2) Scale bars, 10 µm; (A3-D5) Scale bar, 5 µm. (E-J) Quantification of actin cable and Ace-MT features across s10B upon depletion of Dia, Ena, or Villin/Quail. Actin cable density (E) and actin cable length (F) were similarly reduced across the loss of function genotypes. Cortical Ace-MT density (H) was similarly reduced across the genotypes, but cytoplasmic density (G) was only reduced in Villin/Quail mutants. Ace-MT organization (I) was significantly altered only in Villin/Quail mutants. Actin cable and Ace-MT co-alignment (J) was increased across the genotypes. Two-way (E-G, I) or one-way (H) ANOVA with Tukey’s multiple comparison test. (J) Chi-square test for goodness fit. Conditions sharing letters are not statistically different from one another.

### Directly reducing actin filament assembly and bundling results in microtubule network defects

Because MTs are required for actin cable assembly, we next asked if this regulation is bidirectional. We directly perturbed actin filament assembly by reducing Dia or Ena, or actin filament bundling by reducing Villin/Quail, and examined the impact on the Ace-MT network. Reduction of Dia, Ena or Villin/Quail produced significant and virtually identical reductions in actin cable density and growth rate (Fig 7A1-D1, A3-D3, E, F). Reducing actin filament assembly by depleting Ena or Dia did not affect the cytoplasmic Ace-MT network density, organization, or orientation (Fig. 7G, I, Fig. S5A). The fraction of the remaining actin cables that co-aligned with Ace-MTs increased slightly but significantly in Dia and Ena KDs (Fig. 7J, Fig. S5B), but this could be merely the result of a decrease in the total number of cables coupled with a wild type cytoplasmic MT density. Surprisingly, the cortical MT network was significantly depleted in Dia and Ena KDs (Fig. 7A5-C5, H) and this was accompanied by a modest, but significant reduction in cortical Shot in s10B.1 and s10B.2 (Fig. 5I1-J2, O). Shot enrichment on the cables themselves was unaffected (Fig. 5L1-M2, Fig. S4I). Together, these results suggest that loss of cortical MT tethering may be a primary defect. The accumulation of untethered MTs in the cytoplasm may explain the significant increase in perinuclear MT rings in Dia and Ena KDs (Fig. S5C1-2, D).

In contrast, Villin/Quail depletion resulted in a strikingly distinct Ace-MT network. Both cortical and cytoplasmic Ace-MT density was significantly reduced throughout s10B (Fig. 7D2,4, 5, G, H) with no loss of cortical Shot (Fig. 5K1-2, O). In s10B.1-3, the remaining Ace-MTs tended to extend throughout the cytoplasm similar to wild type (Fig. S5E1, E2). We failed to see the shift in Ace-MT orientation bias toward the nucleus at s10B.3 in the Villin/Quail mutants, though this defect was corrected by s10B.4 (Fig. S5F). This cannot be attributed simply to the paucity of cables as in Dia and Ena KDs with a similar cable defect, Ace-MT orientation was unaffected (Fig. S5F). In Villin/Quail mutants, a significantly greater fraction of MTs were in the cortical zone throughout s10B compared to wild type or Dia and Ena KDs (Fig. 7I, Fig. S5A), suggesting that the remaining Ace-MTs are shorter. Furthermore, complete co-alignment between actin cables and the remaining MTs increased through s10B, peaking at s10B.4 significantly higher than any other genotype (Fig. 7J, Fig. S5B). This combination of fewer, shorter, and more co-aligned Ace-MTs created a unique cortically-associated Ace-MT network in Villin/Quail mutants that closely resembled the short, sparse actin cable network in s10B.4 (Fig. 7D1 and D2, D3 and D4).

## Discussion

Here we showed that s10B nurse cells assemble a developmentally regulated, stable, acetylated MT network organized by a cortical *γ*-tubulin/Patronin/Shot based ncMTOC that plays essential roles in the assembly of the actin cable arrays that are necessary for the successful completion of oogenesis (Fig. 8). Centrosomes are often considered the primary MTOCs, but many differentiated cells reassign MTOC function to non-centrosomal sites (Sanchez & Feldman, 2017).

**Figure 8:**
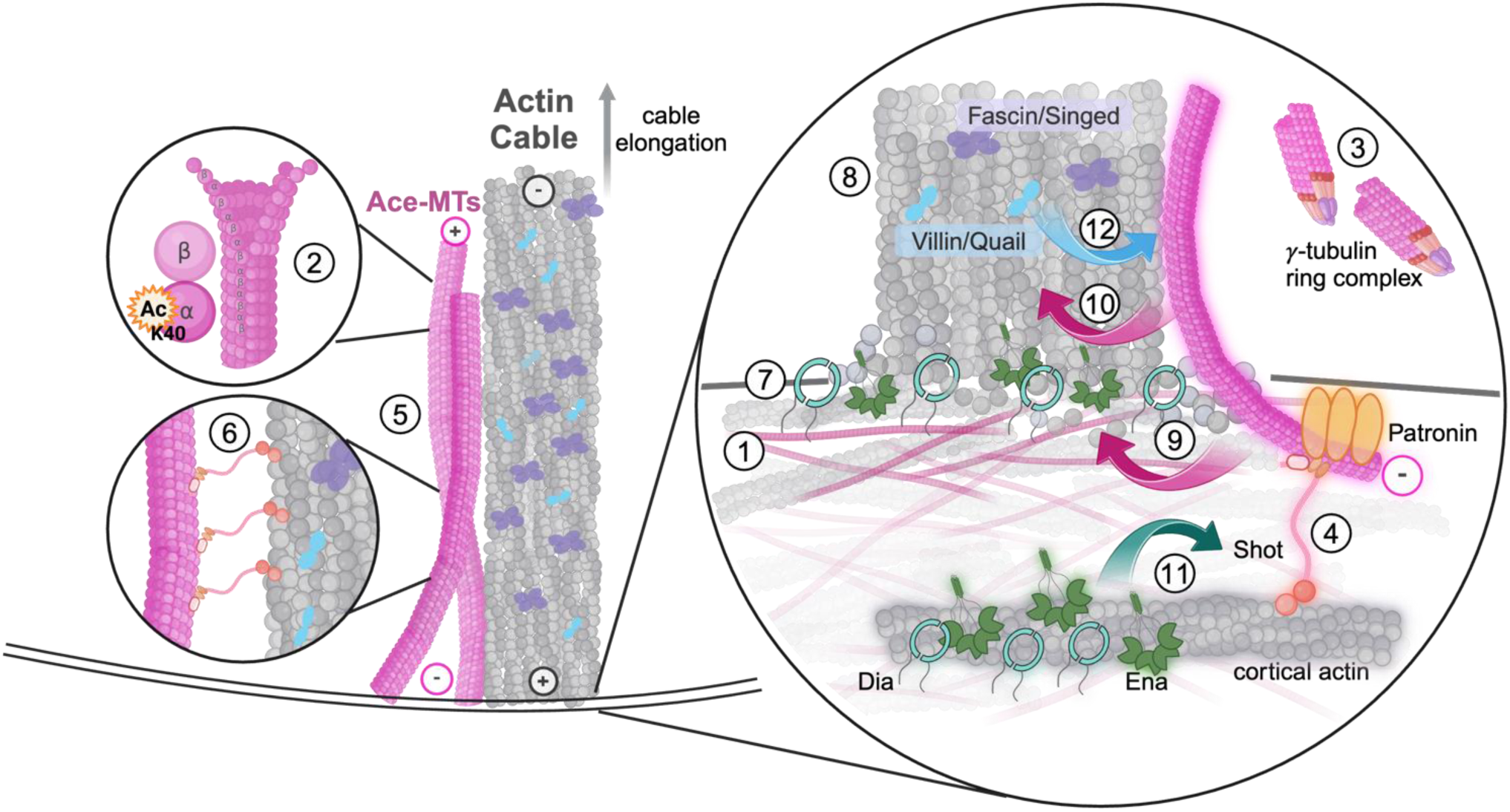
Developmentally regulated crosstalk between actin and Ace-MTs in s10B nurse cells: a cortically organized, acetylated, and stable MT network is required for actin cable assembly, while actin filament assembly and bundling is required to generate and/or maintain the MT network. A stable, acetylated MT network forms a dense meshwork parallel to the cortex (1). Those Ace-MTs (2) are nucleated by *γ*-tubulin that may be acting in the cytoplasm (3). These Ace-MTs are tethered to the actin cortex via Shot and Patronin (4), and extend into the cytoplasm and co-align with actin cables (5). Shot may contribute to actin cable-Ace-MT crosslinking and co-alignment (6). Actin filaments within the cables are nucleated and elongated by cortical Dia and Ena (7), and bundled by Villin/Quail and Fascin/Singed (8). The Ace-MT network is required for actin cable initiation and to promote the cable growth rate. This may be through enhancing the cortical enrichment of Dia and Ena (9). Our data suggest that Ace-MTs regulate actin cable assembly and/or maintenance through other mechanisms as well (10). In turn, Dia and Ena are necessary to maintain the cortical Ace-MT network, possibly through a role in cortical actin and Shot tethering (11). Villin/Quail is required to regulate Ace-MT number, organization and growth (12) potentially through the control of actin cable structure.

At early stages of oogenesis (approx. s2-s5), MTs nucleate from a single MTOC within the oocyte and extend their plus ends through the ring canals into the nurse cells. This provides essential avenues for dynein-mediated transport from the nurse cells into the growing oocyte (Lu et al., 2023; Saxton, 2001; Theurkauf et al., 1992, 1993; Tillery et al., 2018). In s6 and s7, there is a significant reorganization of MTs in both nurse cells (Mische et al., 2007) and oocyte (Steinhauer & Kalderon, 2006; Theurkauf et al., 1992). Through s9, the nurse cell MT network is robust but no MTOC has been identified. At s8-10 in the oocyte, MT minus ends repolarize from the posterior to the anterior membrane, forming an anterior cortical ncMTOC organized by cortical Shot and Patronin foci in the vicinity of cortical *γ*-tubulin foci that were only observed when *γ*-tubulin was overexpressed (Nashchekin et al., 2016). Our data suggests that s10B nurse cells employ a similar Shot/Patronin ncMTOC without apparent cortical *γ*-tubulin (Fig. S4A1-2, B1-2). Cytoplasmic *γ*-TuRCs nucleate MTs in cultured mammalian neurons independent of the MTOC (Yamada & Hayashi, 2019) and could be a source of nucleation here. *γ*-TuRC nucleation can be augmented by Katanin-mediated MT severing of Patronin-stabilized MTs that produces new minus ends for Patronin capture and subsequent plus end growth (Baas & Ahmad, 2013; Lindeboom et al., 2013; Roll-Mecak & Vale, 2006). This may be occurring in the oocyte and follicle cells (Lowe et al., 2014; Nashchekin et al., 2016) and thus MT severing is a good candidate source for new MTs in the nurse cells as well.

MTs in s6-9 nurse cells exhibit dramatically different behaviors from what we observed in s10B (Fig. 3A1, S3E1-F2, supplement video 1, 2; (Lu et al., 2020, 2021, 2022; Theurkauf, 1994). Overall, s6-9 MTs are less stable than what we have observed at s10B and they translocate via Dynein along the cortex and through the ring canals and into the oocyte (Lu et al., 2022). By tracking EB1-GFP and selectively labeling small MT populations, we demonstrated that s10B MTs have enhanced stability: we observed few to no EB1-GFP comets (Fig. 3A2, B, supplemental video 3) and small populations of photoconverted MTs persisted for at least 30 min (Fig 3C1-3, D, supplemental video 4). Furthermore, while the s9 MT network appears minimally acetylated (Fig. S3E1-2), by s10B the entire network is acetylated (Fig. 1C1, Fig. S3G1-2). There is a long-recognized correlation between MT stability and acetylation, but the causal nature of this relationship is not fully understood. Acetylation does not affect tubulin polymerization or depolymerization kinetics *in vitro* (Maruta et al., 1986), but can influence MT structure. Loss of acetylation leads to MTs with fewer protofilaments in *C. elegans* touch-receptor neurons and alters MT organization and morphology (Janke & Magiera, 2020; Topalidou et al., 2012). Additionally, acetylation regulates mechanosensation in *Drosophila* larvae by promoting microtubule stability (Yan et al., 2018). Acetylated MTs are more mechanically stable and resistant to breakage (Xu et al., 2017), but acetylation also appears in less stable structures like the MT network at the cleavage furrow (Danilchik et al., 1998). aTAT1 was not required to maintain the presence of the overall MT network overall in neurons or epithelia (Dan Wei et al., 2018; Shida et al., 2010), but we found that reducing aTAT1 led to dramatic depletion of the total MT network (Fig. 3F1-2). This suggests that acetylation is essential for MT stability and maintenance in the s10B nurse cells (Fig. 8).

The switch from a less stable, largely non-acetylated MT network to a more stable, acetylated one correlates with the end of slow transport of cargo from the nurse cells into the oocyte and the initiation of actin cable assembly in preparation for the rapid expulsion of nurse cell cytoplasm into the oocyte at s11. This suggests that a common signal might trigger both the stability of the MT network and the initiation of actin cable assembly. Prostaglandin (PG) signaling appears to coordinate actin remodeling at s10B: actin cable loss and other defects in PG defective egg chambers were significantly enhanced by reduction of either Ena or Fascin (Groen et al., 2012; Spracklen et al., 2014). Interestingly, loss of PG signaling appeared to cause a reduction in s10B cortical Ena and resulted in premature Ena-dependent actin cable like structures in s9 (Spracklen et al., 2014). Thus, MTs may also be targets of PG signaling, potentially via MT acetylation. aTAT1 can be transcriptionally repressed by the acetyltransferase p300 and activated via phosphorylation by AMP kinase (Mackeh et al., 2014). AMP kinase plays multiple roles in oogenesis (Laws & Drummond-Barbosa, 2016), but a cytoskeletal role has not been explored. Tubulin acetylation can also be increased through the repression of histone deacetylase 6 (HDAC6) that can be regulated through a wide variety of pathways and post-translational modifications (Han et al., 2009; Hook et al., 2002; Williams et al., 2013). Different tubulin isoforms can also contribute to MT stability (Nsamba & Gupta, 2022). In Drosophila, three α-tubulin isotypes, TUB67C, TUB84B, and TUB84D, are essential during oogenesis (Nsamba & Gupta, 2022). Mutations in **α**TUB67C, a maternal specific **α**-tubulin isoform, affect egg size and nurse cell to oocyte ratio (Matthews et al., 1993) – both phenotypes consistent with nurse cell dumping defects.

Depletion of the MT network via *γ*-tubulin, Patronin, Shot, or aTAT1 significantly decreased actin cable density and growth rate, producing defects similar to those we observed with Dia or Ena loss of function (Fig. 6H,I, Fig. 7E,F). This suggested that MT function might be upstream of the localization or activity of both Dia and Ena and indeed we found a significant reduction of cortical Ena and Dia in s10B Shot knockdowns (Fig. 5D2, F2, G-H, Fig. 8). Although significant, this loss of cortical enrichment seemed moderate and unlikely to fully explain the strong loss of actin cables in the absence of MTs. MT may also exert their effects via crosstalking proteins like APC (Adenomatous polyposis coli) (Bahmanyar et al., 2009; Juanes et al., 2019). APC proteins physically interact with actin and the actin cortex (Molinar-Inglis et al., 2018; Moseley et al., 2007; M.-N. Zhou et al., 2011) with Dia, promoting its activity (Breitsprecher et al., 2012; Okada et al., 2010; Webb et al., 2009) and with MTs both directly (Mimori-Kiyosue et al., 2000; Zumbrunn et al., 2001) and via EB1 (Su et al., 1995). EB1 can inhibit APC-mediated actin filament assembly (Juanes et al., 2020; Moseley et al., 2007; Stroud et al., 2014), and with its binding partner CLIP-170 (Cytoplasmic Linker Protein-170) (Gupta et al., 2010) enhances the elongation rate of Dia-mediated actin assembly *in vitro* and promotes Dia activity in cultured cells (Henty-Ridilla et al., 2016; Lewkowicz et al., 2008). Like mammals, flies have two APC proteins and we have previously shown that APC1 and APC2 collaborate with Dia (Jaiswal et al., 2013; Webb et al., 2009), and APC2 regulates cortical actin dynamics in s8 nurse cells (Molinar-Inglis et al., 2018). Interestingly, EB1 and CLIP-170 can also bind actin directly but not simultaneously with MTs (Alberico et al., 2016; Y.-F. O. Wu et al., 2022), although the *in vivo* roles of this function are not yet known. Less is known about interactions between Ena and MTs, but several studies have linked MTs with Abl kinase (Hu et al., 2019; G.-F. Wang et al., 2022), a negative regulator of Ena.

Loss of Dia or Ena had a very similar effect on both the actin and MT cytoskeletons: the reduction in cable number and growth rate (Fig 7E,F, and Logan et al., 2022) and the loss of cortical MTs without a loss of cytoplasmic MTs (Fig. 7G,H) were indistinguishable. The MT defect suggested a loss of cortical MT tethering, and consistent with that we found a significant, though modest reduction in cortical Shot (Fig. 5I2, J2, O, Fig. 8). Actin filament assembly factors like formins and Ena/VASP proteins are cortically localized in a wide variety of organisms and cell types and their loss can have effects on cortical actin and cell-cell adhesions (Baum & Perrimon, 2001; Blake & Gallop, 2023; Chua et al., 2024; Scott et al., 2006). In s9 Drosophila oocytes, thick cortical actin bundles requiring the formin Capu anchor the minus ends of MTs to the cortex (Roth-Johnson et al., 2014; Y. Wang & Riechmann, 2008; Yoo et al., 2015). Thus, the loss of cortical Shot and MT tethering could be an indirect effect of Dia and Ena’s role in cortical actin assembly or dynamics (Fig. 8). Direct effects are also possible; for example, mDia1 directly binds and stabilizes MTs independent of its actin nucleation function in cooperation with EB1 and APC (Bartolini et al., 2008, 2012; Wen et al., 2004).

Cable defects in Villin/Quail mutants were comparable to those in Dia/Ena knockdowns (Fig. 7E,F), however, the remaining actin cables in Villin/Quail mutants are likely the result of bundling by Fascin (Cant et al., 1994, 1998). Villin/Quail and Fascin appear to have different primary roles in actin bundle formation during the production of these actin cables, with Quail acting as a bundle initiator and Fascin as a bundle organizer (Cant et al., 1998). Fascin null mutants rescued by Villin/Quail overexpression produced wild-type-like individual bundles within cables but resulted in disorganized, poorly maintained, and loosely associated cable bundles (Cant et al., 1998). A shift to primarily Fascin-bundled cables may produce the unique MT phenotype observed in Villin/Quail mutants. The increase in actin-MT coalignment (Fig. 7J) and the greater fraction of MTs oriented toward the nucleus are consistent with enhanced crosslinking. The decreased MT length is evidenced by the increased proportion of MTs within the cortical zone. In s10B.3, the few short cables present are predominantly associated with short MTs, while long MTs lack cable associations (Fig. S7E2, E4). In s10B.4, where more cables have initiated and elongated, long MTs are sparse, and most MTs are short and associated with a cable (Fig. 7D1-4). If this reduced MT length results from the shortening of captured long MTs, it suggests that previously stable MTs may shrink to the length of the coaligned cable. These changes could be indirect effects of a change in cable structure, and/or direct effects of altered Fascin availability in those cables or in the cytoplasm. Fascin interacts with and/or regulates Ena (Harker et al., 2019), LIMK (Jayo et al., 2012) and MTs (Villari et al., 2015). The Fascin-MT connection isn’t well understood, but Fascin can bind MTs directly, promote MT dynamics, and may be able to crosslink actin and MTs (Villari et al., 2015). The unique system of actin-MT physical and regulatory interactions we have demonstrated here in the Drosophila nurse cells may provide the opportunity to gain insight into this novel mode of actin-MT crosstalk as well.

## Acknowledgements

Thank you to Simon C. Watkins and the Center for Biologic Imaging (CBI) at the University of Pittsburgh for providing us with invaluable imaging resources. The anti-Dia antibody was a gift from S. Wasserman (UCSD) and M. Baylies (Sloan Kettering Institute). The anti-Patronin antibody was a gift from R. D. Vale (UCSF). The *UASp-Patronin-RNAi* and *Patronin-YFP/CyO* fly lines were a gift from J. Großhans, (Philipps University of Marburg). Many thanks to O. Molinar-Inglis, A. Leslie, G. Logan, and W. Douglas for initiating this work. We thank T. T. Parson and L. A. Raftery (University of Nevada) for sharing their pre-publication work on *Drosophila* centripetal follicle cell migration (Parson et al., 2023), which facilitated our establishment of the s10B staging system. Stocks were obtained from the Bloomington Drosophila Stock Center (NIH P40OD018537). The anti-Ena antibody developed by C. Goodman’s group at University of California, Berkley and anti-Shot mAbRod1 antibody developed by P. A. Kolodziej’s group at Vanderbilt University were obtained from the Developmental Studies Hybridoma Bank, created by the NICHD of the NIH and maintained at The University of Iowa, Department of Biology, Iowa City, IA 52242. FlyBase (release FB2024_05) was used for Drosophila gene information. We also thank all the McCartney lab members for support, discussions, and suggestions. This work was funded by a grant to B. M. McCartney from the National Institutes of Health (R01-GM120378). And by a grant to V.I.Gelfand from the NIH MIRA R35 (National Institute of General Medical Sciences) (2R35GM131752) and the CCBX Research Initiative from the Flatiron Institute/Simons Foundation.

## Methods

### Staging System

Each of the four 10B substages we defined corresponds to a distinct step in the progression of centripetal follicle cell (CFC) migration and the morphological changes of the border cells (BC). Below is a description of their developmental progression and Table 1 defines each substage based on their developmental landmarks (Refer to Table 1 and Fig. S1).

By s9, the anterior follicle cells have flattened to envelop the nurse cells, and the remaining FCs form a columnar secretory epithelium surrounding the oocyte (Parsons et al., 2023; Brigaud et al., 2015). The BCs, a cluster of 6-8 follicle cells, delaminate from the anterior epithelium and migrate between the nurse cells. The BCs migrate towards the oocyte, reaching its dorsal/anterior border by s10A and establishing a stable interface by s10B. Meanwhile during s10B, the CFCs at the nurse cell–oocyte boundary begin migrating inward to envelop the anterior end of the oocyte. Once migration is complete, the BCs and CFCs create a continuous epithelial layer, forming the micropyle by s13 (Miao et al., 2020; Montell et al., 1992).

### Drosophila genetics

Fly stocks and crosses were maintained on standard cornmeal-molasses food supplemented with dry active yeast. Fly stocks were kept at room temperature. To induce expression using Gal4 drivers, crosses were kept at 29℃. The following fly stocks were used in this study: *Oregon-R* (a wild-type control strain), *y^1^w; P{matalpha4-GAL-VP16}67; P{matalpha4-GAL-VP16}15* (*MDD-GAL4*; BDSC_80361), *P{w[+mC]=otu-GAL4::VP16.R}1, w; P{w[+mC]=GAL4-nos.NGT}40; P{w[+mC]=GAL4::VP16-nos.UTR}CG6325[MVD1]* (*MTD-GAL4*; BDSC_31777), *y^1^w; mat atub-Gal4-VP16[V2H] (*BDSC_7062)*; mat atub-Gal4-VP16[V37]* (BDSC_7063), *Ubi-EB1-GFP* (Lu et al., 2021), *UASp-tdEOS2-atub84B* (7M, III, Lu et al., 2013), *UASp-atat-dsRNA* (TRiP.JF03205, attP2, III, BDSC_28777), *UASp-*γ*-tubulin23C-dsRNA* (TRiP.GL01171, attP2, III, BDSC_42799), *UASp-shot-dsRNA* (TRiP.GL01286, attP2, III, BDSC_41858), *UASp-Patronin-dsRNA* (attP2, III, gift from Dr. Jörg Großhans, Philipps University of Marburg), *UASp-dia-dsRNA* (TRiP.HMS00308, attP2, III, BDSC_33424), *UASp-ena-dsRNA* (TRiP.HMS01953, attP2, III, BDSC_39043), *qua[1]cn[1]bw[1]speck[1]/CyO* (II, BDSC_3350), *qua[6-396]/SM1* (II, BDSC_4571), Patronin-YFP/CyO (II, gift from Dr. Jörg Großhans, Philipps University of Marburg).

### Immunostaining

#### Labeling acetylated-MTs (ace-MTs) and actin cables

Flies 1-2 days after eclosion were fed wet yeast paste 24∼48 hours to promote egg production. Ovaries were dissected in freshly prepared Grace’s media (Grace’s medium powder (Sigma, G9771; VWR 95037-608) + 10% Fetal Bovine Serum (FBS) + 7.5% Sodium Bicarbonate (pH 6.0∼6.4), fixed for 20 min in 4% EM-grade paraformaldehyde (PFA) diluted in phosphate-buffered saline (PBS) from an 8% PFA Aqueous Solution (Electron Microscopy Sciences, 157-8) and blocked for 1 h in PBS plus 0.2% Triton X-100 and 4% normal goat serum (NGS) at room temperature. Primary antibody incubation was in PBS plus 0.1% Triton X-100 and 1% NGS with 1:500 Alexa488-Phalloidin (Thermo Fisher Scientific, A12379), 1:350 monoclonal acetylated-⍺-tubulin (Sigma, MABT868; Santa Cruz sc-23950) at 4 ℃ overnight, followed by 1:1000 goat anti-mouse Alexa Fluor™ 568 (Thermo Fisher Scientific A11031) for 3 hr and DAPI (1µg/ml) for 30 min before mounting (Prolong Gold, Invitrogen P36934).

#### Labeling the total MT network

Ovaries were dissected in 1X Brinkley Renaturing Buffer 80 [BRB80, 80 mM piperazine-N,N’-bis(2-ethanesulfonic acid) (PIPES), 1 mM MgCl2, 1 mM EGTA, pH 6.8] followed by a 40 min cytoplasmic extraction: incubation in 1 X BRB80 +1% Triton X-100 for 40 min without agitation. Egg chambers are fixed in 4% EM-grade PFA diluted in BRB80 from an 8% PFA Aqueous Solution + 0.1%Triton X-100 for 20 min at room temperature and blocked for 1 hr at room temperature in 1X PBTB (1X PBS + 0.1% Triton X-100+0.2% BSA). Antibody incubations were in 1X PBTB using 1:500 Alexa568-Phalloidin (Thermo Fisher Scientific A12380), 1:350 FITC-⍺-tubulin (Sigma, F2168), and DAPI (1µg/ml). (Adapted from Lu et al., 2022.)

#### Immunostaining for protein localization

Dissection was performed as previously described for ace-MTs and actin. For *γ*-tubulin and Patronin-YFP localization and Patronin antibody labeling, egg chambers were fixed in 8% EM-grade PFA and 2% Tween for 10 min at room temperature. Samples were blocked with 10% BSA and 0.2% Tween in PBS for 1 hr at room temperature and labeled with primary antibody and 0.2% Tween at 4 ℃ overnight. For Shot, Dia and Ena localization, incubations were performed as described for acetylated MTs and actin cables. Primary antibodies used in this study: 1:500 mouse monoclonal anti-*γ*-tubulin (Sigma T6557), 1:500 chicken polyclonal anti-GFP (Aves Labs GFP-1020), 1:5 mouse monoclonal anti-Shot (shot mAbRod1, DSHB), 1:350 rabbit anti-Patronin (gift from the Vale lab, University of California, San Francisco) and 1:500 rabbit anti-Dia (a gift from the Wasserman lab, University of California San Diego and the Baylies lab, Memorial Sloan Kettering Cancer Center). Samples were incubated in corresponding secondary antibodies (1:1000) for 3 hr at room temperature followed by DAPI (1µg/ml) for 30 min before mounting.

### Nocodazole treatment

For nocodazole treatments to depolymerize MTs, isolated stage 10B egg chambers were transferred into Grace’s media supplemented with 7.5 µg/ml nocodazole (Sigma, #487928) or DMSO and incubated on a shaker at room temperature for 30 min or 60 min. Egg chambers were thoroughly washed with 1X PBS three times before standard fixation for labeling ace-MTs and actin.

### Live Imaging of Drosophila Egg Chambers

Young adult female Drosophila were mated with male flies and fed with active dry yeast paste for 16–18 hours prior to dissection. Ovaries were dissected in Halocarbon Oil 700 (Sigma-Aldrich, Cat# H8898) following established protocols (Lu et al., 2016, Lu et al., 2018).

### Image acquisition

For fixed imaging – Fig. 1F1-4; Fig3. E-H; Fig. 4; Fig. 5; Fig. 6; Fig. 7 A5,B5,C5,D1-5; Fig. S3E1-G2; Fig. S4D1-2; Fig. S5E1-5 samples were imaged on Olympus Fluoview 3000 confocal microscope with a X60/1.42NA. Images were acquired with Fluoview software (FV-10 ASW3.1, Olympus) at 0.4-0.42 µm step size, and a projection of two optical slices was used to show representative images. Fig. 1G1-4; Fig. S4 (excluding 4D1-2) were imaged with an Evolve EMCCD, others with a Prime95B CMOS camera (Photometrics) on a Zeiss Axiovert 200 M microscope with an X-Light V2 spinning disc scan head (Crest Optics). Fluorophores were excited with a Celesta Light Engine (Lumencor). Images were acquired with MetaMorph software at 0.2 µm step size, and a projection of five optical slices was used to show representative images. Cortical MT images are projections of ten 0.4 µm optical slices.

For live imaging – Freshly dissected samples were imaged using a Nikon W1 spinning disk confocal microscope (Yokogawa CSU with a 50 µm pinhole size), equipped with a Hamamatsu ORCA-Fusion Digital CMOS camera and a 40× 1.25 N.A. silicone oil lens, controlled by Nikon Elements software.

### Live imaging of EB1 comets

Homozygous *ubi-EB1-GFP* flies (Lu et al., 2015, Lu et al., 2023) were used. Images were acquired at a frame rate of one frame every 2 seconds over a 2-minute period.

### Photoconversion of *tdEOS2-⍺-tubulin*

F1 progeny expressing *tdEOS2-αtub* (7M, III) (Lu et al., 2013, Winding et al., 2016), driven by double maternal αtub-Gal4 drivers (*mat αtub-Gal4[V2H]; mat αtub-Gal4[V37]*), were used. Photoconversion from green to red was performed locally in nurse cells using 395 nm light from an Aura Lumencor light source at 100% power, controlled by a Mightex Polygon DMD Illuminator, for 30 seconds. Photoconverted microtubule images were acquired at frame rates of one frame every 10 seconds for a total of 10 minutes or one frame every 30 seconds for a total of 30 minutes.

### Quantification, statistical and image analysis

All experiments were replicated two to three times. n refers to either number of egg chambers or cytoskeletal components specified. Measurements and basic image processing were performed with Fiji (Image J v2.14; National Institution of Health) and Adobe Photoshop (v25.9.1). All statistical analyses and graphs were generated with Prism 10 (GraphPad). Figures were prepared using Microsoft Powerpoint. Schematics and models were created with BioRender.

#### Categorization of EB1-GFP comet dynamics

The classification of Ubi-EB1-GFP egg chambers based on the presence of EB1-GFP puncta in nurse cells is as follows: Egg chambers with 30% to 100% of frames showing active EB1-GFP puncta in nurse cells were categorized as "frequent comets”. Those with fewer than 30% of frames showing active EB1-GFP puncta were categorized as "few comets". Those with no frames showing active EB1-GFP puncta were categorized as "no comets”. n=4-12 egg chambers per substage.

#### Categorization of photoconverted MTs in live egg chambers

Photoconverted egg chambers from s10A to s10B.4 were categorized based on the traceability of labeled MTs. Egg chambers were classified as having trackable MTs if the majority of labeled MTs could be observed throughout the imaging period (either 10 or 30 minutes) or as non-trackable if the majority could not be followed to the end of imaging. n=4-13 egg chambers per sub/stage.

#### Quantification of actin cable properties

For quantifying actin cable density, the total number of actin cables was measured over five spans of nurse cell borders totaling minimally 120 µm in length. Analysis performed on images of wild-type (all 10B substages), MTD>OR (all 10B substages), and knockdown/mutants (s10B.3 and s10B.4), taken on the Prime95B CMOS camera or Evolve EMCCD used a projection of five

0.2 µm optical slices. For images of wild-type (all 10B substages), MTD>OR (all 10B substages), and knockdown/mutants (10B.3 and 10B.4) taken on Olympus Fluoview 3000 confocal microscope used a projection of two 0.4µm optical slices. For analysis performed on all knockdown and mutants in stages 10B.1 and 10B.2, we measured cables in nurse cells across the z-plane (40∼50 µm z stack) to ensure complete coverage of the nurse cell. n=5-12 egg chambers per substage. For quantifying actin cable growth rate, we measured the length of 10 to 20 actin cables (if available) on at least three different nurse cell borders in each egg chamber. For substages and knockdown/mutant samples with fewer cables, we measured all the cables present. n=5-10 egg chambers per substage.

#### Quantification of MT properties

For quantifying cytoplasmic MT density, images taken on the Prime95B CMOS camera or Evolve EMCCD used a projection of five 0.2 µm optical slices. For images taken on Olympus Fluoview 3000, we used a projection of two 0.4 µm optical slices. Cytoplasmic MT density in nurse cells was calculated by dividing the total number of intersection points (yellow asterisks) by the total cytoplasmic area, calculated from the number of 4.25 μm^2^ regions (yellow squares) covering the cytoplasm (Fig. S2A1-2). n=5-12 egg chambers per substage. For cortical MT density, projections of ten (0.4 µm step size; Olympus Fluoview 3000) or twenty (0.2 µm step size; Prime95B CMOS camera or Evolve EMCCD) optical slices were generated and cortical MT intersections were counted in over 100 µm^2^ of phalloidin-labeled cortex per egg chamber. n =4-12 egg chambers per substage.

For quantifying Ace-MT organization, we divided the cytoplasm into the “nuclear zone” and the “cortical zone”. This was achieved by marking the halfway point between the nucleus and the cortex and connecting the marks to create borders parallel to the cortex, equally dividing the cytoplasm into halves. We then counted the Ace-MT intersections within each zone to assess the Ace-MT density distribution (Fig. S2B), n=5-10 egg chambers per substage. For quantifying orientation, Ace-MTs were analyzed if they met the following criteria: longer than 3 µm, had no more than three intersections or discontinuities, and had clearly identifiable start and/or end points. A ruler tool (Fig. S2C1, yellow), divided into 45° intervals, was applied to determine where Ace-MTs fell within the 90° range. The center line was aligned perpendicular to the cortex, adjusting the cortex parallel to the ends of the Ace-MTs (Fig S2C2-3, yellow dotted line). In wild-type, Dia and Ena knockdowns, 20-50 distinct Ace-MTs were analyzed per egg chamber. For other knockdowns and Villin/Quail mutants with cytoplasmic Ace-MT density defects, all Ace-MTs meeting these criteria were analyzed. n=5-10 egg chambers per substage.

#### Quantification of actin and Ace-MT co-alignment

For quantifying actin and Ace-MT co-alignment, we selected actin cables where its full length was clearly visible. If an Ace-MT completely overlaid the entire length of an actin cable, it was categorized as "completely co-aligned”. If an Ace-MT overlaid only part of the actin cable, it was categorized as "partially co-aligned”. If there was no Ace-MT at any point along the entire length of the actin cable, it was categorized as "not co-aligned”. 20-60 actin cables were analyzed per egg chamber. n=5-10 egg chambers per substage.

#### Quantification of fluorescence intensity

Fluorescence intensity for Dia, Ena, and Shot was quantified in fixed egg chambers by measuring the intensity across 2-3 segments, each spanning 6-10 μm from the nurse cell border to the cytoplasm, or 3-8 μm from the actin cable to the cytoplasm. The relative cortical or cable-to-cytoplasm intensity ratio was calculated for each segment. Cortical and cable regions were identified based on colocalization with actin. Measurements are taken from at least two different nurse cells within a single egg chamber, with a total of 5-10 egg chambers analyzed per substage per genotype.

