## Supplementary figures for "Developmentally regulated actin-microtubule crosstalk in Drosophila oogenesis"

#### Supplemental Material Legends

Supplemental **Video 1: EB1-GFP in s2-3 egg chambers.** Live imaging of EB1-GFP was used to track growing MT plus ends. In s2-3 egg chambers, abundant EB1 comets can be observed running parallel to the cortex and throughout the cytoplasm, as well as in the follicle cells. Scale bar, 5 µm. (Corresponds to Fig. 3A1-A2). Images acquired every 2 sec over a period of 2 minutes.

Supplemental **Video 2: EB1-GFP in s9 egg chambers.** In s9 egg chambers, abundant EB1 comets can be observed in nurse cells running parallel to the cortex and throughout the cytoplasm, as well as in the follicle cells. Scale bar, 30 µm. Images acquired every 2 sec over a period of 2 minutes.

Supplemental **Video 3: EB1-GFP in s10B.3 egg chambers.** In s10B.3, the frequency of EB1 comets in nurse cells decreased significantly. The few comets present were predominantly near and parallel to the cortex. In contrast, EB1-GFP comets are active and abundant in follicle cells. Scale bar, 30 µm. (Corresponds to Fig. 3A1-A2). Images acquired every 2 sec over a period of 2 minutes.

Supplemental **Video 4**: **Photoconversion of tdEOS2-αtub.** In stage 10B.2, a subpopulation of cytoplasmic MTs labeled with photoconverted tdEOS2-αtub. Scale bar, 5 µm. (Corresponds to Fig. 3C1-C3). Each frame is 30 sec apart, imaged over a 30 minute period.

Supplemental Figure 1: **Using the migration of centripetal follicle cells (CFCs) and border cells (BCs) to define substages in s10B Drosophila oogenesis.** (A1-A4) Schematic of stage 10B subdivided into four substages (10B.1 to 10B.4) defined by the progressive movement of centripetal FCs (green) at the nurse cell-oocyte boundary and the visibility of border cells. The extent of CFC migration is indicated as follows – 50% (yellow dashed lines) and 80% (red dashed lines). Refer to Table 1 for a detailed definition of s10A to s11 centripetal follicle cell migration and border cell morphology in relation to the progression of actin cable assembly. (B1-G1) Representative fixed images s10A to s11 wild-type egg chambers stained with Phalloidin and DAPI. (B1-G1) Scale bars, 40 µm; (B2-G5) Scale bars, 10 µm. (B2-G2) Border cells. (B3-G4) Centripetal follicle cells. (B5-G5) Nurse cell. (H-J) Wild type actin cables initiate and elongate in s10B and actin cable density is not correlated with cytoplasmic Ace-MT density. (H) Quantification of actin cable density in wild-type nurse cells across s10B. (I) Actin cable length across s10B. (H)(I) Significance is compared to the previous substage. One-way ANOVA, ****$p<0.0001$, Tukey's post-hoc analysis. (J) Actin cable density and cytoplasmic Ace-MT density correlation across s10B. Pearson correlation, $P=0.6263$. $n=10$ per substage.

Supplemental Figure 2: **Quantification methods for MT density, organization, orientation and actin coalignment.** (A1-A2) Cytoplasmic MT density in nurse cells was calculated by dividing the total number of intersection points (yellow asterisks) by the total cytoplasmic area, calculated from the number of 4.25x4.25 µm regions (yellow squares) covering the cytoplasm). Scale bar, (A1) 10 µm, (A2) 1 µm. (B) MT organization was quantified by marking the halfway point between the nucleus and cortex. These marks were connected to create borders parallel to the cortex, dividing the cytoplasm into two equal halves: the “nuclear zone” (adjacent the nucleus) and the “cortical zone” (adjacent to the cortex). MT density was then calculated for each zone. Scale bar, (B) 10 µm. (C1-C3) MT orientation was quantified by selecting MTs that meet the specified criteria (see Methods). A ruler tool (C1, yellow), divided into 45º intervals, was applied to determine where MTs fell within the 90º range. The center line was aligned perpendicular to the cortex, adjusting the cortex parallel to the ends of the MTs (yellow dotted line). For example, (C2) MT #4 falls within the “perpendicular to nucleus (45-90º)” range, while (C3) MT #13 falls within the “parallel to nucleus (0-45º)” range. Scale bar, (C1) 10 µm, (C2-C3) 2 µm. (D1-F3) Actin-MT co-alignment was categorized into three groups: (D1-D3) Completely co-aligned—where a fully visible actin cable is entirely aligned with a fully visible Ace-MT; (E1-E3) Partially co-aligned—where only part of the actin cable is aligned with an Ace-MT; and (F1-F3) Not co-aligned—where no portion of the actin cable is aligned with an Ace-MT. Scale bar, (D1-F3) 2 µm.

Supplemental Figure 3: **Acetylated MTs form an intertwined cytoplasmic network extending from a dense cortical meshwork at s10B.** Wild-type examples of (A1-A3) Ace-MTs with one end embedded in the cortex (yellow and white empty arrowheads) and the other extending into the cytoplasm (yellow and white filled arrowheads). The white arrowheads in A1 mark the same Ace-MT as in A2. (B) An example of an Ace-MT that lies parallel to the cortex (empty arrowheads), with the other end extending outward into the cytoplasm (filled arrowhead). MTs associated with (C1) or passing through (C2) ring canals between nurse cells (arrowheads). Prominent perinuclear (PN) MT rings were observed occasionally in s10B.4 (D2), but most nurse cells did not have PN rings (D1). (D3) Categorization of PN ring phenotypes in wild-type s10B.4 nurse cells (n=31). (E1-G2) Prior to s10B, the nurse cell cytoplasmic MT network is largely non-acetylated in both s9 (E1-E2) and s10A (F1-F2). Though we did see Ace-MTs at the cortex in these stages. (A1, B, D1-D2, E-G1) Scale bars, 10 µm. (E-G2) Scale bars, 5 µm. (A2-A3, C1-C2) Scale bars, 2 µm.

Supplemental Figure 4: ***γ*-tubulin, Patronin, and Shot localization in s10B nurse cells.** (A1-C2) γ-Tubulin was not significantly enriched at any specific subcellular location. We observed occasional weak enrichment at the cortex (B1-B2, arrowhead) or perinuclear region (C1-C2, arrowhead). (C3) Centrosomes were clearly labeled in early stage egg chambers. (A1-A2) Scale bar, 10 µm; (B1-C3) Scale bar, 5 µm. (D1-E2) Patronin is strongly enriched at the cortex (D1-D2) with occasional enrichment in the cytoplasm near the nucleus (E1-E2; arrowhead) Scale bar, 5 µm. (F1-F2) Patronin localization in *MDD>shot dsRNA*. Patronin is no longer cortical upon Shot depletion. Scale bar, 10 µm (G1-G2) Patronin is not required for Shot localization at the cortex or along cables. Scale bar, 10 µm. (H) Comparison of Shot enrichment along cables in s10B.3 and s10B.4 wild type, *MDD>dia dsRNA*, *MTD>ena dsRNA*, and *qua[1]/qua[6-396]* nurse cells. Shot cable enrichment was unaffected. One-way ANOVA, n=5-11 (as plotted). Tukey’s multiple comparison test.

Supplemental Figure 5: **Direct perturbation of actin filament assembly and bundling leads to a range of MT network defects.** (A) Quantification of MT organization in control, *MDD>dia dsRNA*, *MTD>ena dsRNA*, and *qua[1]/qua[6-396]* in s10B.2 to s10B.4. Villin/Quail mutants shifted to a significantly more cortically organized cytoplasmic MT network across s10B. Conditions sharing letters are not statistically different from one another. (B) Quantification of actin cable and Ace-MT co-alignment in s10B.2 to s10B.4. The only significance observed was in s10B.4 Villin/Quail mutants (****$p<0.0001$). All other genotypes/substages do not differ significantly. (C1-C2) There was a significant increase in the presence of perinuclear MT rings in Dia and Ena knockdowns. (D) Categorization of PN ring phenotypes in genotypes indicated. (E1-E5) s10B.3 actin and MT network in Villin/Quail mutants. Long MTs that are present lack cable associations. (E1, E3) Low and high magnification of actin cables. Scale bar, 10 μm. (E4) Cytoplasmic and (E5) cortical MTs. Scale bar, 5 μm. (F) Quantification of MT orientation across s10B. Villin/Quail mutants fail to show the MT orientation shift at s10B.3. This appears to be independent of cable defects as orientation remained unaffected in Dia and Ena knockdowns with comparable cable defects. Two-way (A) or one-way (F) ANOVA, with Tukey's multiple comparison test. n=5-10 per substage. (B, D) Chi-square test for goodness fit.

### Table 1

| Stage | Centripetal follicle cells (CFCs) | Border cells | Actin cable assembly |
|---|---|---|---|
| 10A | CFCs are positioned at the anterior edge of the columnar FCs. They are oriented at a slightly oblique angle to the oocyte cortex but have not yet begun to stretch. (Fig.1A1, S1B3-4) | BCs have reached the nurse cell/oocyte boundary. The BC nuclei, cluster delineation, and outlines of individual follicle cells are clearly visible with phalloidin-actin. (Fig. S1B2) | No actin cable initiation. (Fig. S1B5) |
| 10B.1 | The leading CFCs at the anterior edge have begun to stretch apically. The subsequent CFCs have started to thin out and are more laterally compressed along the apical-basal axis. (Fig.S1A1, C3-4) | Border cell cluster remains strongly present. Border cell nuclei, cluster delineation, and individual follicle cell outline within, are clearly visualizable. (Fig. S1C2) | Light cable initiation (cable density average<0.1cables/µm) with minimal growth (average length < 2µm). (Fig. 1D1, S1C5) |
| 10B.2 | CFCs have elongated and begun migrating, with one or both sides covering no more than 50% of the distance from the nurse cell/oocyte border to the center. (Fig.S1A2, D3-4) | The BC nuclei, cluster boundaries, and individual follicle cell outlines are generally clear. With Phalloidin, 15-20% of cases show less sharply defined follicle cell outlines compared to stage 10B.1. (Fig. S1D2) | Increase in cable initiation (average=0.2 cables/µm) and modest degree of cable growth (average length 4.6 µm). (Fig. 1D2, S1D5) |
| 10B.3 | CFCs have migrated significantly toward the center. Migration falls between 50% and 80% of the anterior border, with one or both sides reaching this range. (Fig.S1A3, E3-4) | BC cluster outline is still visible with phalloidin-actin staining. In 50% of cases, the cluster appears smaller, making it harder to distinguish individual follicle cell outlines within the border cells. (Fig. S1E2) | Robust cable density (average=0.38cables/µm) and elongation (average length 8 µm). (Fig. 1D3, S1E5) |
| 10B.4 | Both sides have migrated at least 80% of the anterior border toward the center. CFCs are either close to or completed migration. Columnar FCs are thinnest in stage 10B. (Fig.S1A4, F3-4) | The border cell outline is least distinct during stage 10B. In 90% of cases, we were unable to visualize the cell cluster with phalloidin-actin. (Fig. S1F2) | Cable arrays reached peak density (average 0.53 cables/µm), with many cables elongating to contact with the nuclei (average length 13 µm). (Fig. 1D4, S1F5) |
| 11 | CFCs have completed migration. (Fig.1A3, G3-4) | Border cells clusters are rarely seen (<5%). with phalloidin-actin. (Fig. S1G2) | Actin cables reposition nuclei as cytoplasmic dumping occurs, with oocyte length exceeding that of the nurse cells. (Fig. S1G1, 5) |

### Figure S1

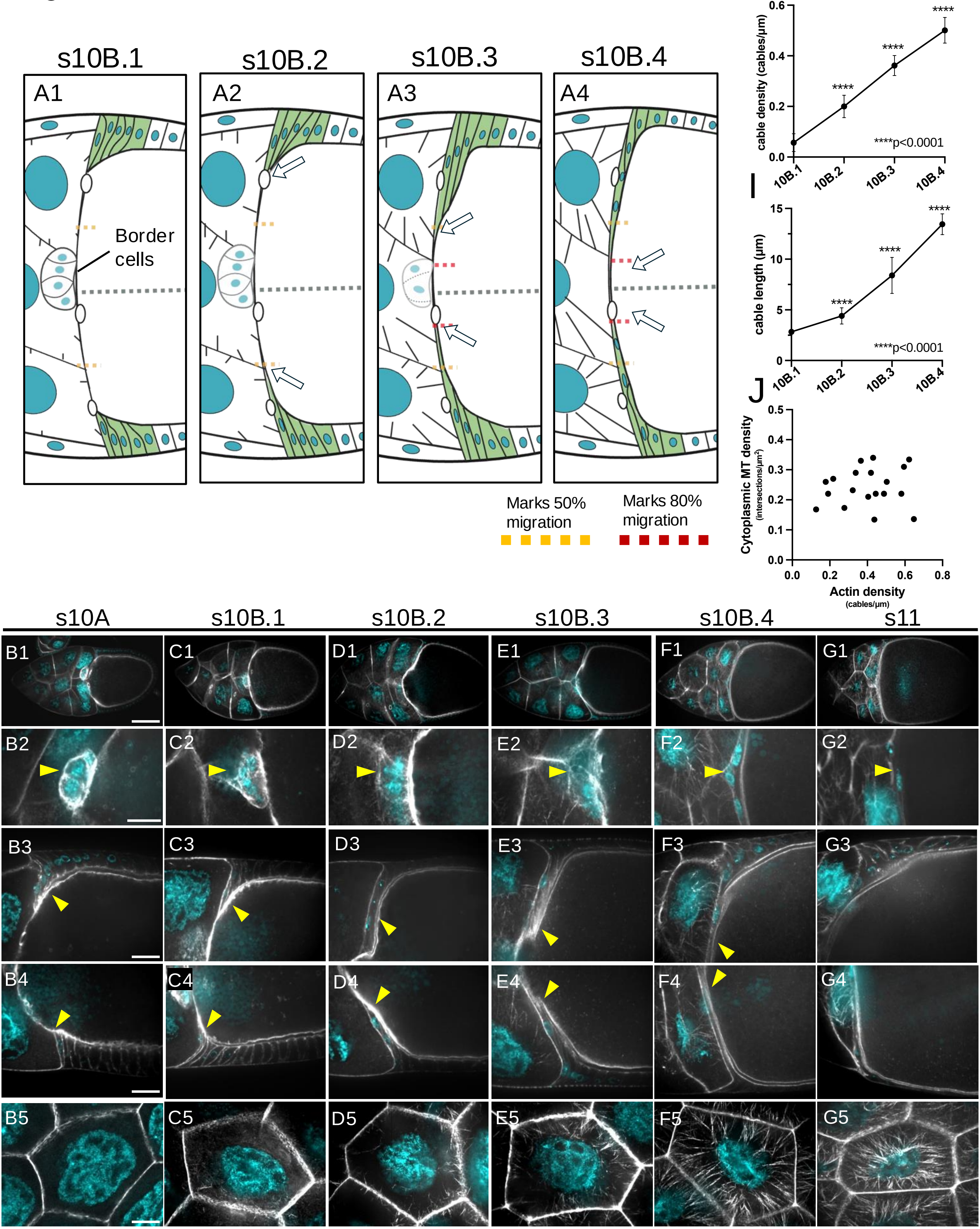

s10B.1
s10B.2
s10B.3
s10B.4
A1
A2
A3
A4
Border cells
Marks 50% migration
Marks 80% migration
H
cable density (cables/µm)
****p<0.0001
10B.1
10B.2
10B.3
10B.4
I
cable length (µm)
****p<0.0001
J
Cytoplasmic MT density (intersections/µm²)
Actin density (cables/µm)
s10A
s10B.1
s10B.2
s10B.3
s10B.4
s11
B1
C1
D1
E1
F1
G1
B2
C2
D2
E2
F2
G2
B3
C3
D3
E3
F3
G3
B4
C4
D4
E4
F4
G4
B5
C5
D5
E5
F5
G5

### Figure S2

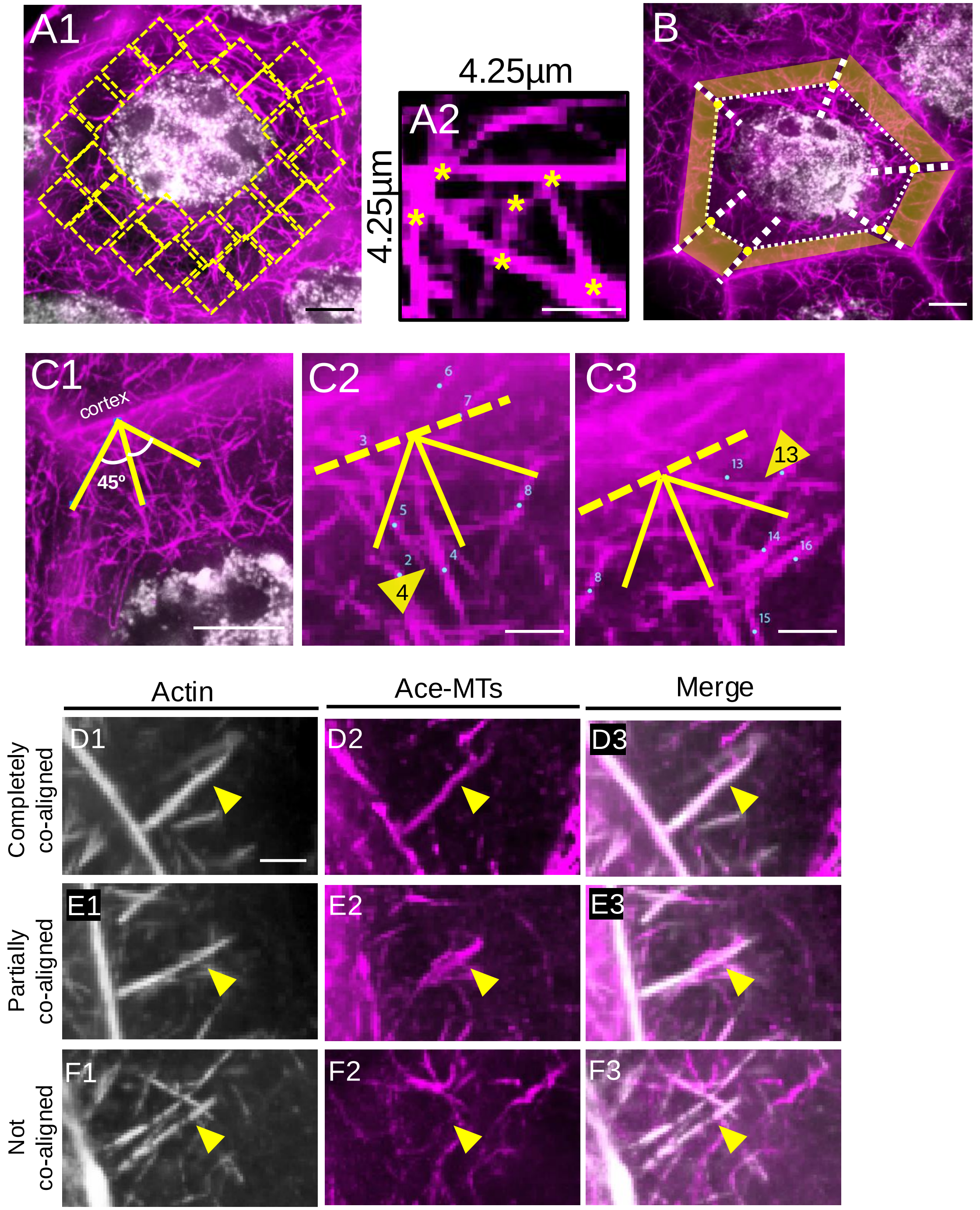
A1
4.25µm
A2
4.25µm
B
C1
cortex
45º
C2
3
6
7
8
5
2
4
C3
13
14
16
8
15
Actin
Ace-MTs
Merge
Completely co-aligned
Partially co-aligned
Not co-aligned
D1
D2
D3
E1
E2
E3
F1
F2
F3

Figure S3

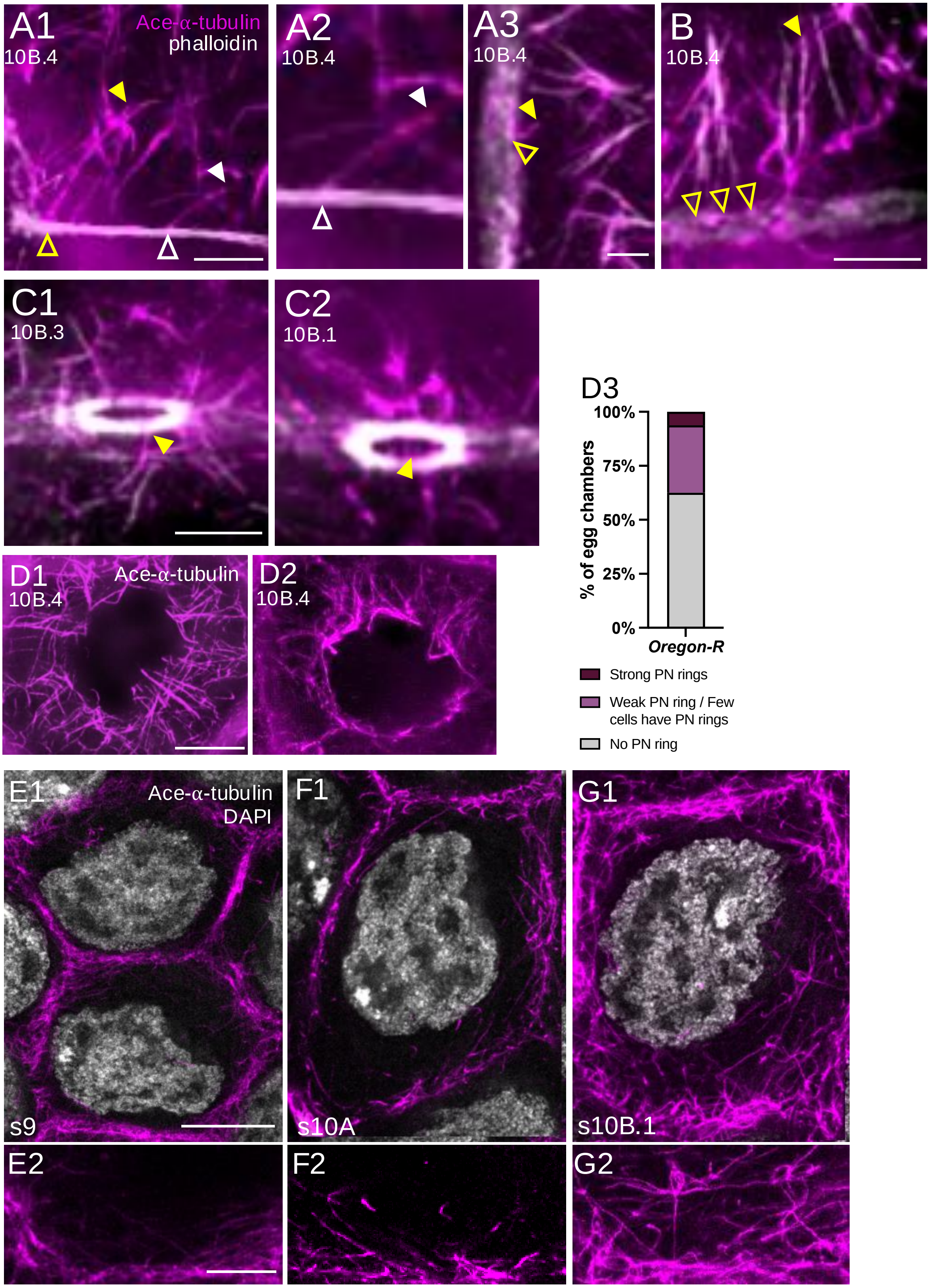

A1
Ace-α-tubulin
phalloidin
10B.4
A2
10B.4
A3
10B.4
B
10B.4
C1
10B.3
C2
10B.1
D3
100%
75%
50%
25%
0%
% of egg chambers
Oregon-R
Strong PN rings
Weak PN ring / Few cells have PN rings
No PN ring
D1
Ace-α-tubulin
10B.4
D2
10B.4
E1
Ace-α-tubulin
DAPI
s9
F1
s10A
G1
s10B.1
E2
F2
G2

### Figure S4

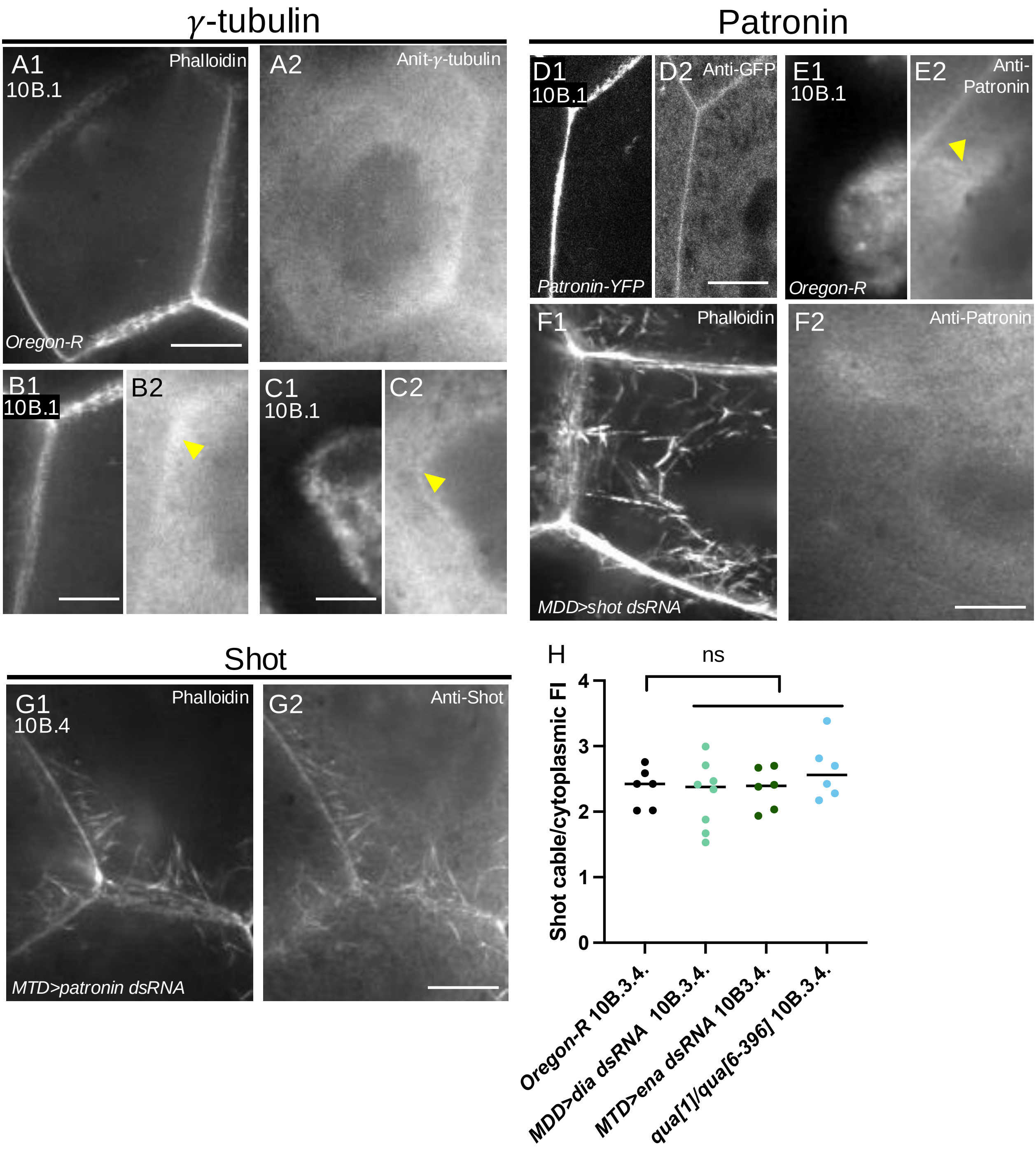
γ-tubulin
Patronin
A1
10B.1
Phalloidin
A2
Anit-γ-tubulin
Oregon-R
B1
10B.1
B2
C1
10B.1
C2
D1
10B.1
D2
Anti-GFP
Patronin-YFP
E1
10B.1
Oregon-R
E2
Anti-Patronin
F1
Phalloidin
MDD>shot dsRNA
F2
Anti-Patronin
Shot
G1
10B.4
Phalloidin
MTD>patronin dsRNA
G2
Anti-Shot
H
ns
Shot cable/cytoplasmic FI
0
1
2
3
4
Oregon-R 10B.3.4.
MDD>dia dsRNA 10B.3.4.
MTD>ena dsRNA 10B3.4.
qua[1]/qua[6-396] 10B.3.4.

### Figure S4

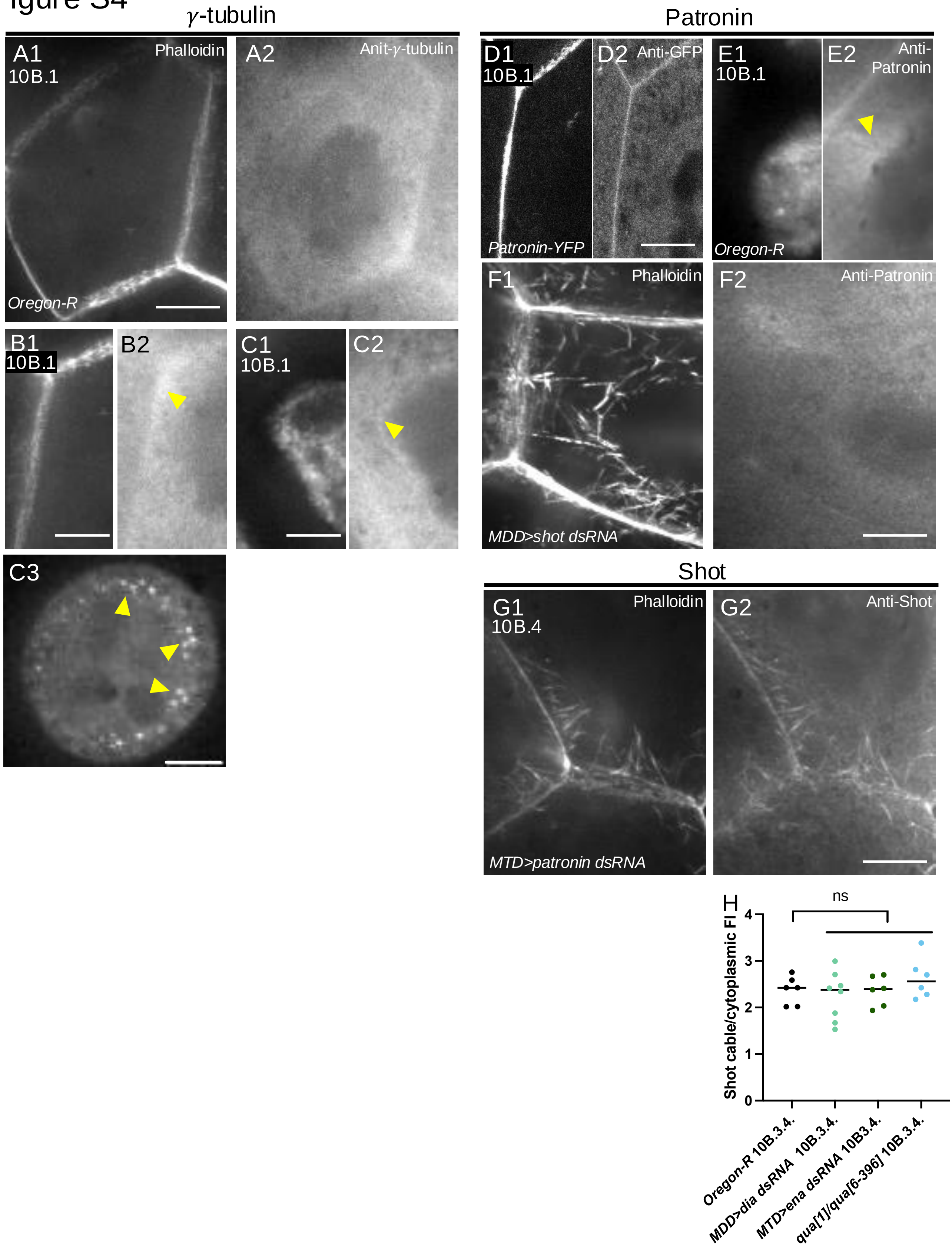

γ-tubulin
Patronin
A1
10B.1
Phalloidin
Oregon-R
A2
Anit-γ-tubulin
D1
10B.1
Patronin-YFP
D2
Anti-GFP
E1
10B.1
Oregon-R
E2
Anti-Patronin
B1
10B.1
B2
C1
10B.1
C2
F1
Phalloidin
MDD>shot dsRNA
F2
Anti-Patronin
C3
Shot
G1
10B.4
Phalloidin
MTD>patronin dsRNA
G2
Anti-Shot
H
ns
Shot cable/cytoplasmic FI
0
1
2
3
4
Oregon-R 10B.3.4.
MDD>dia dsRNA 10B.3.4.
MTD>ena dsRNA 10B3.4.
qua[1]/qua[6-396] 10B.3.4.

### Figure S5

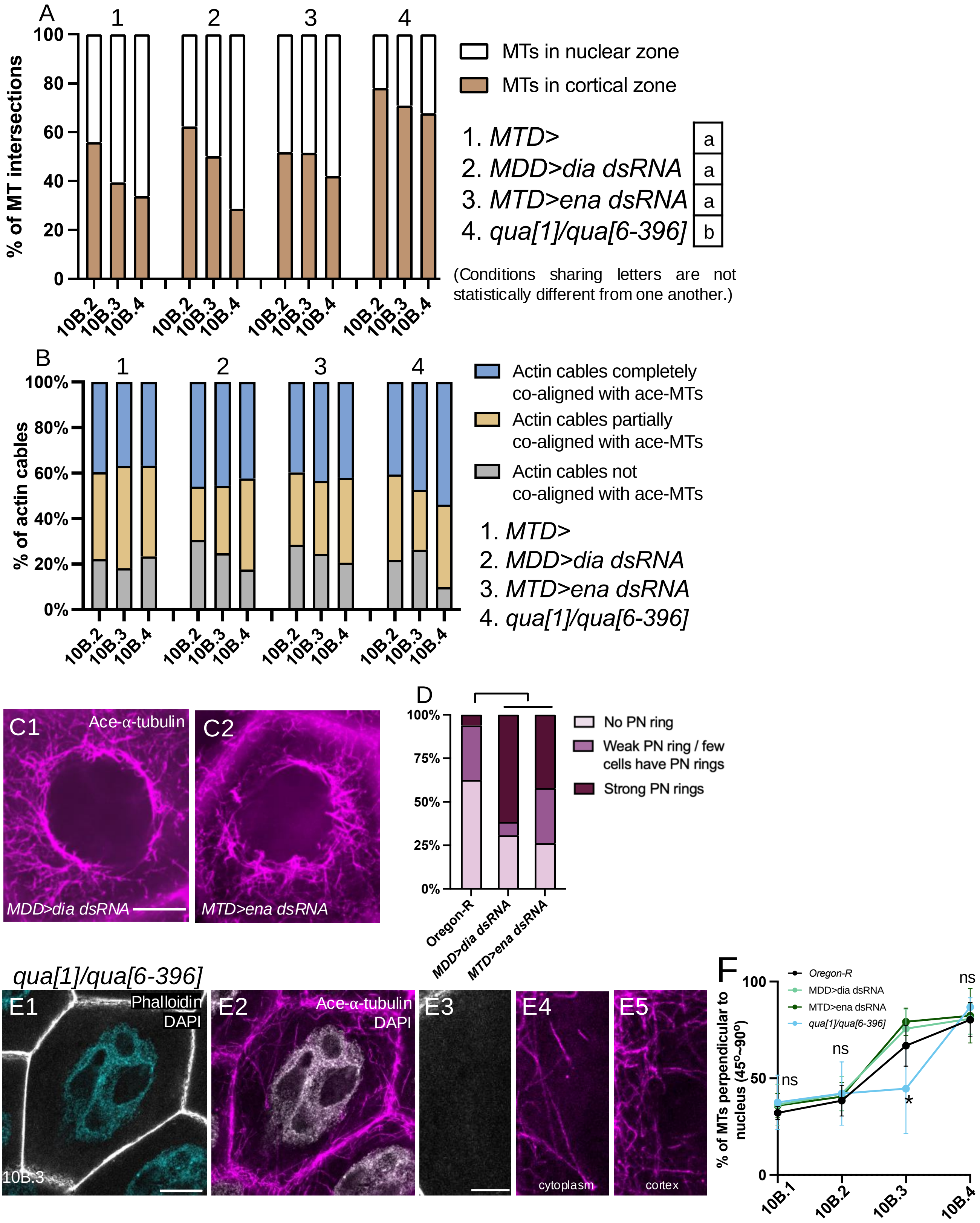

A
1
2
3
4
% of MT intersections
100
80
60
40
20
0
10B.2
10B.3
10B.4
MTs in nuclear zone
MTs in cortical zone
1. MTD> a
2. MDD>dia dsRNA a
3. MTD>ena dsRNA a
4. qua[1]/qua[6-396] b
(Conditions sharing letters are not statistically different from one another.)
B
% of actin cables
100%
80%
60%
40%
20%
0%
Actin cables completely co-aligned with ace-MTs
Actin cables partially co-aligned with ace-MTs
Actin cables not co-aligned with ace-MTs
1. MTD>
2. MDD>dia dsRNA
3. MTD>ena dsRNA
4. qua[1]/qua[6-396]
C1
Ace-α-tubulin
MDD>dia dsRNA
C2
MTD>ena dsRNA
D
100%
75%
50%
25%
0%
Oregon-R
MDD>dia dsRNA
MTD>ena dsRNA
No PN ring
Weak PN ring / few cells have PN rings
Strong PN rings
qua[1]/qua[6-396]
E1
Phalloidin
DAPI
10B.3
E2
Ace-α-tubulin
DAPI
E3
E4
cytoplasm
E5
cortex
F
Oregon-R
MDD>dia dsRNA
MTD>ena dsRNA
qua[1]/qua[6-396]
% of MTs perpendicular to nucleus (45°~90°)
100
50
0
ns
ns
*
ns
10B.1
10B.2
10B.3
10B.4
